# Subcellular localization of mutant P23H rhodopsin in an RFP fusion knockin mouse model of retinitis pigmentosa

**DOI:** 10.1101/2021.10.11.463949

**Authors:** Michael A. Robichaux, Vy Nguyen, Fung Chan, Lavanya Kailasam, Feng He, John H. Wilson, Theodore G. Wensel

## Abstract

The P23H mutation in rhodopsin (Rho), the visual pigment protein in rod photoreceptor neurons, is the most common genetic cause of autosomal dominant retinitis pigmentosa (adRP), a retinal disease that causes blindness. Despite multiple studies in animal models, the subcellular details of the fate of misfolded mutant Rho in rod photoreceptors have not been completely defined. We generated a new mouse model of adRP, in which the P23H-Rho mutant allele is fused to the fluorescent protein Tag-RFP-T (P23HhRhoRFP). In heterozygotes, outer segments formed, and WT rhodopsin was properly localized there, but mutant P23H-Rho protein was specifically mislocalized in the inner segments of rods. Despite this cellular phenotype, the P23HhRhoRFP heterozygous mice exhibited only slowly progressing retinal degeneration; in ERG recordings, scotopic a-wave amplitudes were reduced by 24% and 26% at 30 days and 90 days respectively, and the corresponding scotopic b-waves by 18% and 24%. Outer nuclear layer thickness was still 80% of WT at 90 days, but at 364 days had declined to 40% of WT. Transmission electron microscopy revealed greatly expanded membrane lamellae in the inner segment, and by fluorescence imaging, we determined that the mislocalized P23HhRhoRFP was contained in greatly expanded endoplasmic reticulum (ER) membranes. TUNEL staining revealed a slow pace of cell death involving chromosomal endonucleolytic degradation. Quantification of mRNA for markers of ER stress and the unfolded protein response revealed little or no increases in levels of messages encoding the proteins BiP, CHOP, ATF6, XBP1, PERK, Eif2α and Derlin-1, but a decreased level of total *Rhodopsin* (mouse + human) mRNA levels. The decline in the rate of cell death after an initial burst suggests that P23HhRhoRFP mutant rods undergo an adaptative process that prolongs survival despite gross P23HhRhoRFP protein accumulation in the ER. Because of its slowly progressing nature, and easy visualization of the mutant protein, the P23H-Rho-RFP mouse may represent a useful tool for the future study of the pathology and treatment of P23H-Rho and adRP.

## Introduction

Retinitis pigmentosa (RP) is a hereditary disease of photoreceptor neurons of the retina that causes night blindness, retinal degeneration and, eventually, complete blindness. RP accounts for half of the known cases of all inherited retinal disease (1), affecting 1:4000 in the US (2). Rod and cone photoreceptors, the light-sensing cells in the vertebrate retina are polarized neurons with a specialized outer segment (OS) sensory cilium. The OS is the site of phototransduction, the pathway initiated by the photopigment rhodopsin (Rho) in rods photoreceptors and the cone opsins in cones.

Rho is a prototypical G protein-coupled receptor (GPCR) that is densely packaged as an integral transmembrane protein into the lipid bilayers of membrane discs that fill the rod OS cilium. Each mouse OS contains ∼800 discs, and Rho comprises >90% of the total membrane protein content in the OS (3, 4). Every day ∼10% of the OS membrane mass in mammalian rods is renewed, as the distal OS discs are shed and engulfed by retina pigment epithelium (RPE) phagocytosis (5, 6), and new discs are generated at the base of the OS (7, 8). Thus, in each mouse rod photoreceptor ∼2.4 million Rho molecules must be synthesized daily in the inner segment (IS), the biosynthetic compartment of rods, to be delivered to the OS cilium to sustain this renewal (3). This trafficking load – including Rho, other visual proteins, and lipids - must pass through a thin, 300 nm connecting cilium (CC) bridge located between the IS and OS. The CC has a cylindrical 9 microtubule doublet cytoskeletal core similar to the structure of the transition zone of other primary cilia (9–11), that extends into the OS and nucleates from a pair of basal body (BB) centrioles located at the distal end of the IS in rods.

The correct localization of Rho throughout the highly specialized compartments of rods is necessary to satisfy the high trafficking rate of Rho molecules in rods, and maintain homeostasis and cell survival (12). Thus, rods are susceptible to genetic mutations to Rho itself, which are the leading cause of autosomal dominant RP (adRP) (13). One point mutation that encodes a proline-to-histidine change at codon 23 (P23H) in the N-terminus of Rho is the most common single mutation cause of adRP in North America (14, 15).

P23H-Rho is a misfolding mutation that causes mislocalization of P23H-Rho protein in cell culture from the plasma membrane to dense endoplasmic reticulum (ER) cytoplasmic aggregates (16–18). The deleterious effect of the P23H-Rho mutation has been extensively studied in rods in a wide range of animal models, where it causes rod cell death and retinal degeneration with different degrees of severity (e.g., (19–21). In transgenic P23H-Rho frog rods, the mutant *Xenopus*-P23H-Rho protein is specifically retained and mislocalized in the IS ER (22), while transgenic bovine-P23H-Rho protein in frog rods caused light-induced vesiculation in the IS (23). A zebrafish model expressing mouse-P23H-Rho caused photoreceptor degeneration, abnormal rod OS formation and P23H-Rho protein mislocalization throughout the malformed rods in adult fish retina (24).

Transgenic and knockin P23H-Rho mouse models have variable retinal degeneration rates that coincide with inconsistent photoreceptor cell phenotypes across models (25–33). Transgenic P23H-Rho mice generated in a Rho null background have severely dysmorphic rods, in which mutant P23H-Rho is mislocalized in endoplasmic reticulum (ER) surrounding the nucleus (30). In contrast, a knockin mouse model, featuring a mouse-P23H-Rho knockin allele, had dramatic mutant P23H-Rho protein degradation and no detectable ER mislocalization or accumulation phenotype (28). Rod degeneration in the P23H-Rho knockin heterozygotes was fairly rapid (the rod population was reduced by 50% before P40 (34)) but even more severe in homozygotes. Notably, the undegraded P23H-Rho protein in the P23H-Rho knockin rods normally localized to the OS and caused abnormal OS disc formation (28, 34). One challenge for characterizing the fate of misfolded P23H-Rho and testing its effects on different therapeutic approaches, is the difficulty in distinguishing WT rhodopsin from a variant that differs by one amino acid residue. We previously generated a P23H-Rho-GFP fusion knockin mouse to easily visualize the mutant protein in rods, and we observed a gross mislocalization of mutant P23H-Rho-GFP in the IS (35). Furthermore, we found that the mutant fusion protein was largely degraded; however, we did not characterize any subcellular phenotypes in single rods on a nanoscopic level in the P23H-Rho-GFP mouse (21)

For a more in-depth characterization we have generated a new P23H-Rho mouse model in order to study mutant P23H-Rho protein localization and dynamics, and to serve as a platform for testing gene-based and other therapies. The latter goal is best achieved with a model that has a relatively slow rate of retinal degeneration in line with the slowly progressive course of human vision loss in adRP, which would permit long-term studies. In addition, a relatively non-perturbing tag greatly benefits the testing of therapeutic interventions and strategies.

In this study we introduce a new knockin mouse model of P23H-Rho adRP, in which the mutant P23H-Rho protein is fused to a photostable Tag-RFP-T (36) fusion tag, which we have used to study the mislocalization of the protein. We have also studied the morphological and functional changes that accompany a slowly progressing retinal degeneration

## Results

### Generation of the P23H-hRho-RFP knockin mouse

We designed a new mouse line in which we introduced into the mouse *Rho* locus, the human rhodopsin gene (introns and exons) encoding both the P23H mutation in the first exon and a fusion to Tag-RFP-T at the C-terminus. Tag-RFP-T is a red fluorescent protein (excitation maximum = 555 nm, emission maximum = 584 nm) based on a naturally occurring anemone protein engineered for both bright fluorescence and enhanced photostability (36). Notably, Tag-RFP-T fluorescence is compatible with GFP fluorescence for multiplex detection, and it has been used for multiple applications since its design, including as a fluorescent fusion marker (37–39).

In addition, we added an additional 1D4 signaling sequence to the carboxy-terminus of the P23H-hRho + Tag-RFP-T fusion allele (Fig 1A-B). Our rationale was that the C-terminal RFP fusion tag in the expressed P23H-hRho-Tag-RFP-T (hereafter P23HhRhoRFP) protein may inhibit the endogenous Rho 1D4 sequence leading to artifacts not attributable to the P23H mutation. The additional 1D4 sequence was also shown to be necessary for normal trafficking of transgenic Rho-Dendra fusion protein in *Xenopus laevis* rods (40).

**Figure 1.**
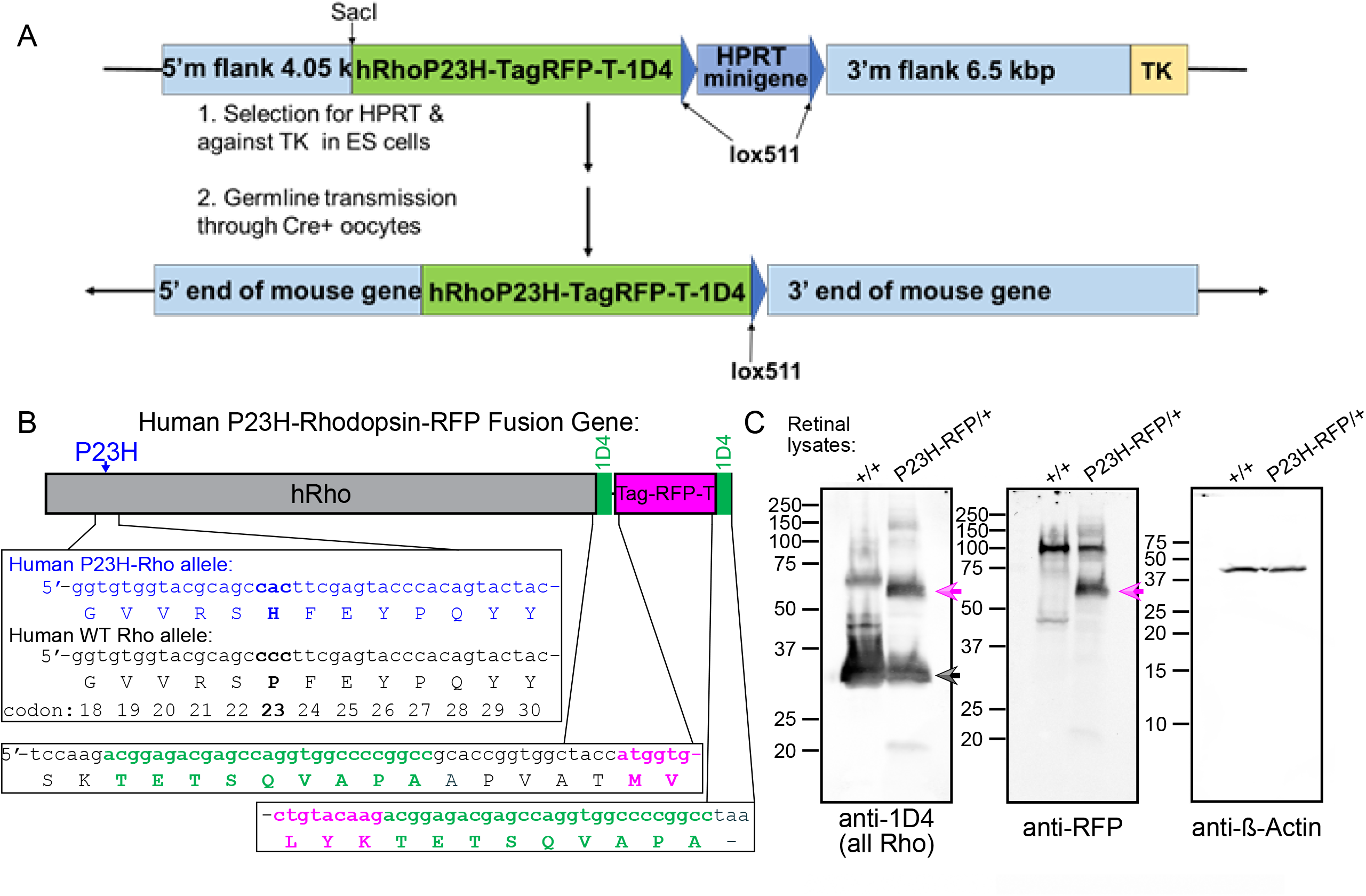
Construction and validation of the P23H-hRho-TagRFP-T knockin mouse. (A) Targeting construct used in embryonic stem cells and resulting gene structure after germline transmission. (B) A map of the knock-in human P23H-rhodopsin-RFP fusion gene. A portion of the sequence of exon 1 from the P23H-Rho allele in the fusion gene (blue) is aligned to the same human wild-type (WT) Rho allele sequence (black). The mutated codon 23 is in **bold** font. The transition sequence from the 1D4 terminal signal sequence to the Tag-RFP-T with a short linker sequence is shown in the middle sequence panel and the C-terminal sequence with the extra 1D4 epitope sequence appended to the end Tag-RFP-T prior to the stop codon. (C) Western blot confirmation of the P23HhRhoRFP fusion protein expression in P23H-RFP/+ heterozygous retinas. Retinal lysates are from a wild-type (+/+) mouse, age P22, and a P23H-RFP/+ mouse, age 45; 100 µg of total protein from each lysate was loaded onto SDS-PAGE gels. Blot membranes were probed with either of the following antibodies: anti-1D4, anti-RFP or anti-beta (ß)-actin (a loading control). Blot scans for the protein ladder were used to mark molecular weight sizes (in kilodaltons, kDa) on the left of each blot image. The ∼65 kDa P23HhRhoRFP fusion protein band is present in the P23H-RFP/+ lane in both anti-1D4 and anti-RFP blot scans (magenta arrows). The monomeric mouse Rho protein band is in both lanes in the anti-1D4 blot scan (black arrow). Higher MW species are formed by rhodopsin multimerization.

The P23H-hRho-TagRFP knock-in mice were generated the same way we previously generated P23H-hRho-GFP knock-in mice (35), by gene targeting in the HPRT^−^ embryonic stem (ES) cell line AB2.2 123, which was derived from mouse strain 129SvEv, essentially as described previously (41, 42). We introduced the P23H mutation into the targeting vector by site-directed mutagenesis (QuikChange^Ⓡ^, Stratagene). An ISceI recognition site was engineered into the middle of the first intron in the rhodopsin gene at position 1340 from the start of translation, but it was not used in the experiments described here. The Darwin Transgenic Core Facility, Baylor College of Medicine, electroporated ES cells, selected for HPRT^+^TK^−^ cells and injected correctly targeted ES cells into blastocysts from albino C57BL/6-Tyr^c-Brd^ mice (43). Founder mice carrying the HPRT-P23H-hRho-TagRFP allele were crossed to GDF-9-iCre mice (44) to remove the HPRT minigene. P23H-hRho-TagRFP (hereafter “P23H-RFP” in reference to the knockin allele) mice were extensively backcrossed to C57BL/6 mice. We validated that the knockin was successful by sequencing genomic DNA from the knockin mouse. We verified expression of the P23HhRhoRFP fusion by fluorescence microscopy of retinas and by immunoblotting (Fig. 1, Fig. 2).

**Figure 2.**
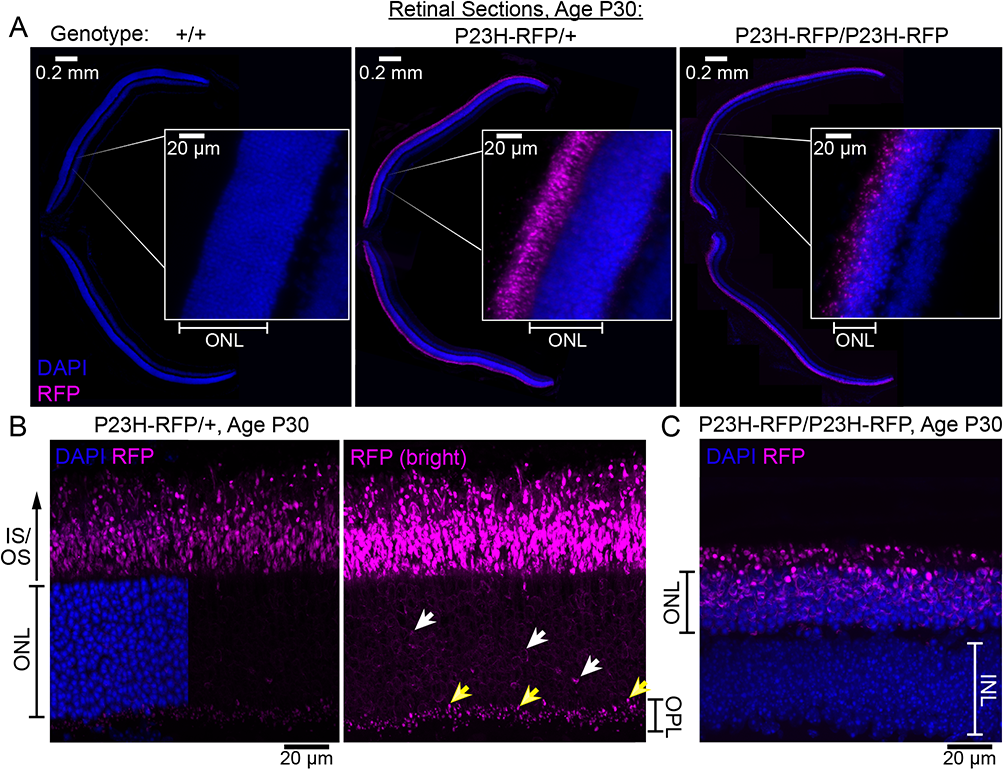
Localization of RFP fluorescence in the P23H-hRho-RFP knockin mouse retina. (A) Widefield fluorescence images of mouse retinal cryosections from age-matched P30 wild-type (+/+), heterozygous (P23H-RFP/+) and homozygous knockin mice (P23H-RFP/P23H-RFP). Sections were counterstained with DAPI to label nuclei in the retina (blue). RFP fluorescence from the mutant P23HhRhoRFP fusion protein is shown in magenta. Magnified regions from each section are insets, and the outer nuclear layer (ONL) of each retina, which is the location of the DAPI+ photoreceptor nuclei, is demarcated. (B) Confocal z-projection image of a retinal cryosection from a P30 heterozygous P23H-RFP/+ mouse. The brightest RFP signal is in the outer photoreceptor layers, the inner segment and outer segment region (IS/OS), where P23HhRhoRFP protein is localized in aggregates. In the same image, with the gain of the RFP signal raised to saturation, additional P23HhRhoRFP protein is observed in the rod photoreceptor synapses within the outer plexiform layer (yellow arrows). To a lesser degree, P23HhRhoRFP is also localized in the ONL, in the cytoplasm surrounding the photoreceptor nuclei (white arrows). (C) Confocal z-projection through a retinal cryosection from a P30 P23H-RFP/P23H-RFP homozygote. The width of the DAPI-positive ONL is thinner compared to the heterozygote. In the homozygous retina, mutant P23HhRhoRFP fusion protein is also more prominently localized in the ONL compared to the heterozygote. INL = inner nuclear layer.

We used P23H-RFP/+ heterozygous mouse retinas to examine the expression of the mutant fusion protein alongside the WT mouse Rho protein from the wild-type allele (Fig 1B). Both WT mouse Rho protein and the product of the knockin allele were detected in P23H-RFP/+ retinal lysates by probing with anti-1D4 antibody. The P23HhRhoRFP fusion protein was detected as a strong ∼65 kDa band specifically in P23H-RFP/+ lysates using both anti-1D4 and anti-RFP antibodies, verifying robust expression.

### P23HhRhoRFP protein was mislocalized in rod photoreceptor neurons

We next tested the fluorescence pattern of the P23HhRhoRFP fusion protein in the retinas of both P23H-RFP/+ heterozygous and P23H-RFP/P23H-RFP homozygous mice. The RFP fluorescence is evident in the outer photoreceptor layers of retinal sections from both heterozygotes and homozygotes at age P30 (Fig 2A). In a confocal z-projection of a retinal section from a P30 P23H-RFP/+ heterozygote, P23HhRhoRFP is most prominently located in brightly fluorescent puncta or “aggregates” within the regions of the inner segments and outer segments of photoreceptors (Fig 2B). Less prominent but visible in this same section is P23H-Rho-RFP fluorescence at the photoreceptor synapses of the outer plexiform layer (OPL) and in the cytoplasm surrounding the photoreceptor nuclei of the outer nuclear layer (ONL) (Fig 2B). Compared to both WT and heterozygotes, the ONL of P30 homozygotes was noticeably thinner based on DAPI+ nuclei staining (Fig 2C). P23HhRhoRFP aggregates were also clearly visible in the ONL of the homozygous retina.

We crossed our new P23H-RFP mouse line with wild-type hRho-GFP mouse lines for dual fluorescent tag multiplex imaging. This approach allowed us to investigate the subcellular dynamics of the mutant P23HhRhoRFP protein relative to hRho-GFP without the P23H mutation. In addition to the hRho-GFP fusion mouse that we previously reported (41), we also generated a new line with an additional 1D4 signal sequence added to the end of the EGFP sequence (“hRho-GFP-1D4”). As heterozygotes, we could not discriminate any phenotypic difference between these GFP fusion mice.

Both GFP fusion lines were crossed to our new P23H-RFP knockin mouse. In confocal images of retinal sections from an adult hRho-GFP-1D4/P23H-RFP heterozygote we observed a drastic localization difference between hRho-GFP-1D4, which correctly populated the rod photoreceptor OS cilia, and the P23HhRhoRFP aggregates, which were prominently mislocalized in the rod inner segment layer (Fig 3A). We examined retinal sections from this same hRho-GFP-1D4/P23H-RFP heterozygous mouse line with structured illumination microscopy (SIM) superresolution imaging and observed a clear segregation of P23HhRhoRFP from the hRho-GFP-1D4 in the OS cilia (Fig 3B). Interestingly, in retinal sections from the other heterozygous hRho-GFP/P23H-RFP mice, we found evidence of hRho-GFP mislocalization in the same region as the P23HhRhoRFP aggregates in what appear to be distinct but interwoven membrane compartments (Fig 3C). This result suggests that the added C-terminal 1D4 sequence prevents hRho-GFP from being mislocalized with P23HhRhoRFP in the GFP/RFP heterozygous animals.

**Figure 3.**
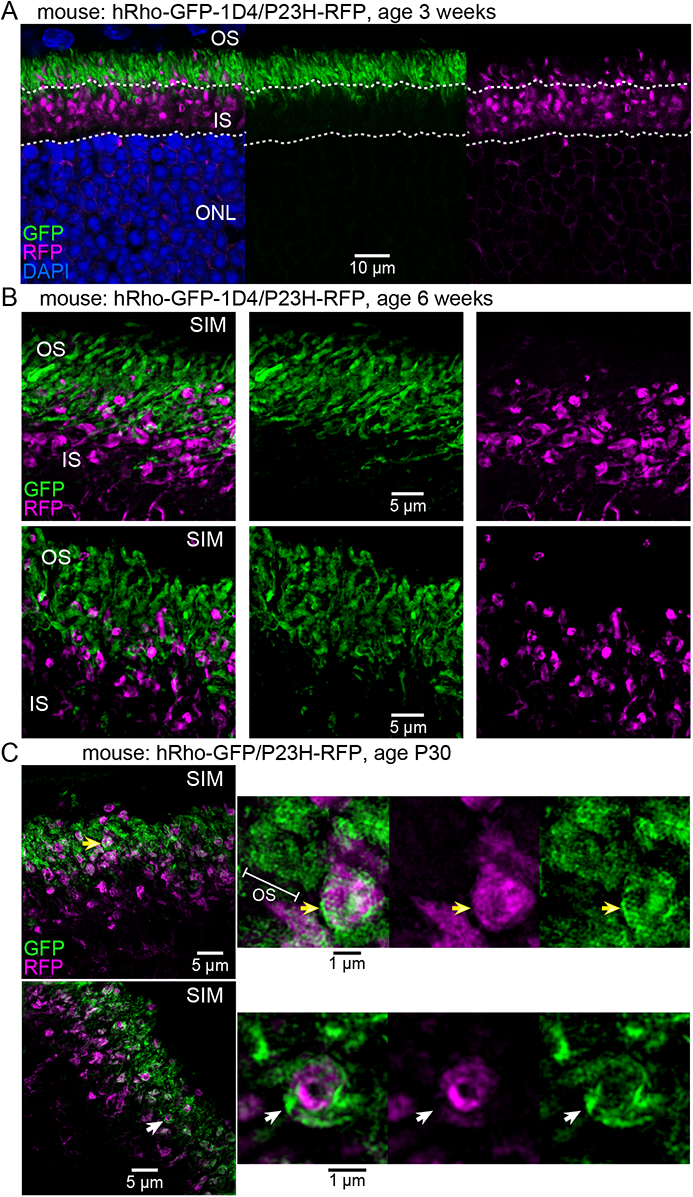
P23HhRhoRFP+ fusion aggregates are localized in the inner segments of rod photoreceptor neurons. (A) In a confocal z-projection image of a retinal section from the hRho-GFP-1D4/P23H-RFP heterozygous mouse retina, at 3 weeks of age, the wild-type hRho-GFP fusion (green), which has an additional C-terminal 1D4 signal sequence, is correctly localized to the outer segments (OS) of rod neurons. In this retina, RFP+ aggregates containing mutant P23HhRhoRFP protein (magenta) are primarily located in the inner segments (IS) of rods and almost entirely segregated from the GFP+ OS layer. White dotted lines in the figure demarcate the OS/IS boundary and the IS/outer nuclear layer (ONL) boundary. (B) In SIM micrographs of retinal sections from the same heterozygous mouse line at age 6 weeks, GFP+ OS and RFP+ IS aggregates remain segregated with no apparent co-localization. (C) In SIM images of retinal sections from an alternate GFP/RFP heterozygote at age P30, in which the wild-type hRho-GFP fusion does not have an additional 1D4 signal peptide, the GFP fluorescence is not exclusively located in the OS layer, but rather is partially mislocalized with P23H-hRho-RFP. In a magnified example, hRho-GFP is co-localized around and within a P23HhRhoRFP aggregate (yellow arrows). In another magnified example, hRho-GFP is wrapped around an P23HhRhoRFP aggregate (white arrow).

To visualize the location of the P23HhRhoRFP inner segment aggregates relative to the connecting cilium (CC) and basal body (BB) we used centrin as an antibody marker for the CC and BB (45, 46) in retinal sections from P23H-RFP/+ mice and imaged by SIM. At age P14, we found many examples of P23HhRhoRFP aggregates that were located just proximal to the BB (Fig 4A). We observed this same sub-BB localization of the RFP aggregates in retinal sections from age P30 P23H-RFP/+ mice (Fig 4B), with some examples of P23HhRhoRFP fluorescence overlapping into the BB region. In retinas from both ages, the RFP fluorescence pattern within these P23HhRhoRFP clumps was discontinuous with dark patches. Morphologically, the P23HhRhoRFP fluorescent aggregates in P14 retina had a compact and defined elliptical shape compared to the aggregates in the P30 retina which appeared more elongated and less structured.

**Figure 4.**
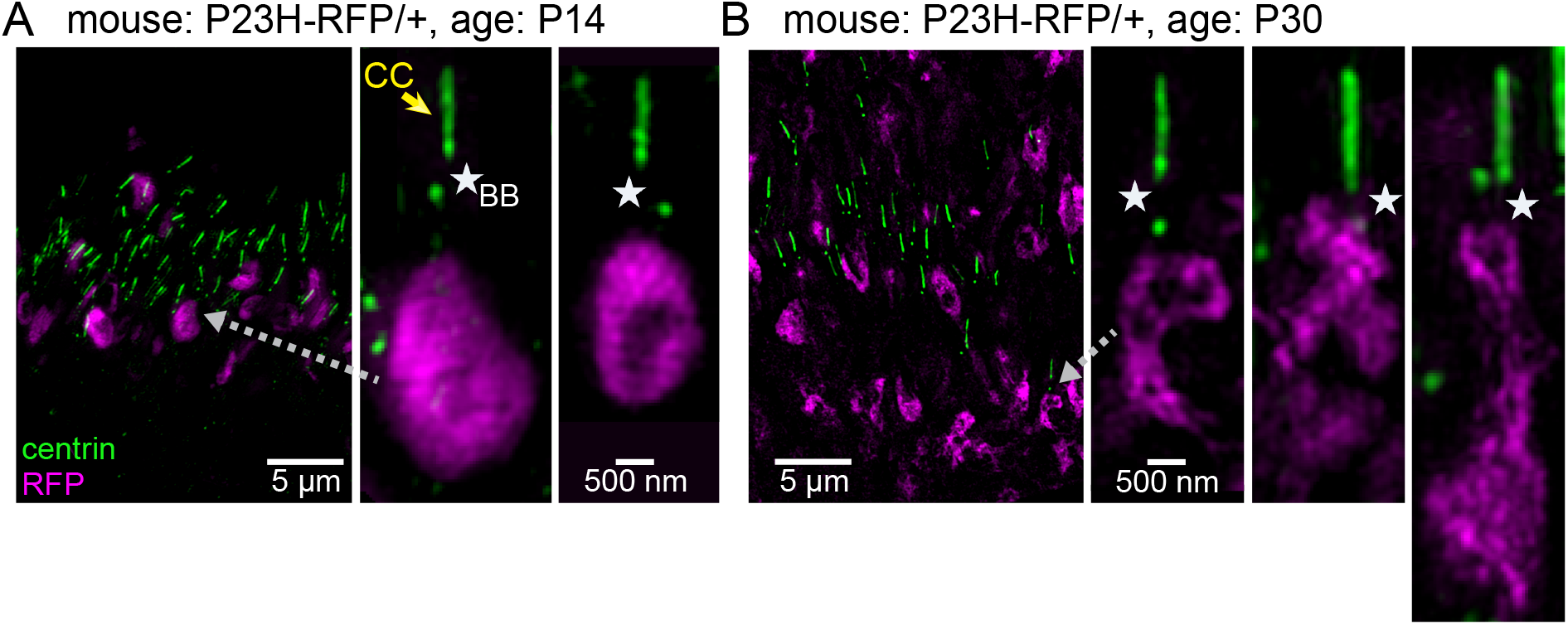
Mutant P23H-hRho-RFP inner segment aggregates are localized near the basal body in rod neurons. (A) In a SIM z-projection image of a retinal section from the P23H-RFP/+ mouse at age P14, centrin immunolabeling (green) marks the location of rod connecting cilium (CC) and basal body (BB) relative to the P23HhRhoRFP fluorescent aggregates (magenta). In magnified views, single RFP+ inner segment aggregates are located just proximal to the CC (yellow arrow) and BB (white stars). The BB region is demarcated by the centrin+ mother centriole at the proximal end of the CC and the daughter centriole, which is a separated centrin+ puncta beneath the mother centriole. (B) A SIM image of a retinal section from the P23H-RFP/+ mouse at age P30 also with centrin immunolabeling. In magnified views, the RFP+ aggregates at P30 are still generally located proximal to the BB (white stars), with some examples of RFP overlapping with the BB region. The organization of the RFP+ aggregates in these P30 rods are less compact and more reticulated compared to the examples from age P14 (see examples in A).

Taken together, fluorescence images of P23H-RFP retinas at different ages demonstrate that the P23HhRhoRFP mutant fusion protein was excluded from the OS and mislocalized within aggregates near the BB in the IS and to a lesser degree in the ONL and the photoreceptor synapses. This result suggests that the mutant fusion protein is accumulating in a trafficking stage just prior to the BB and integration into the cilia, and thus it fails to be properly transported to the OS.

### P23H-RFP/+ mice have mild and gradual retinal degeneration

Next, we tested the effect of the mutant P23HhRhoRFP fusion on retinal health by measuring the rates of retinal degeneration in both P23H-RFP/+ heterozygous and P23H-RFP/P23H-RFP homozygous mice. We measured the thickness of the ONL, in which the photoreceptor nuclei were stained for 4′,6-diamidino-2-phenylindole (DAPI) fluorescence, in retinal sections from both genotypes and in wild-type (+/+) control retinas from mice at multiple timepoints. Overall, the P23H-RFP/+ heterozygous retinas degenerated more slowly over time compared to the homozygous retinas as measured by ONL thickness (Fig 5A-B). Our analysis covered the full retina – peripheral to central – to test for any region-specific degeneration caused by the mutant P23HhRhoRFP fusion protein.

**Figure 5.**
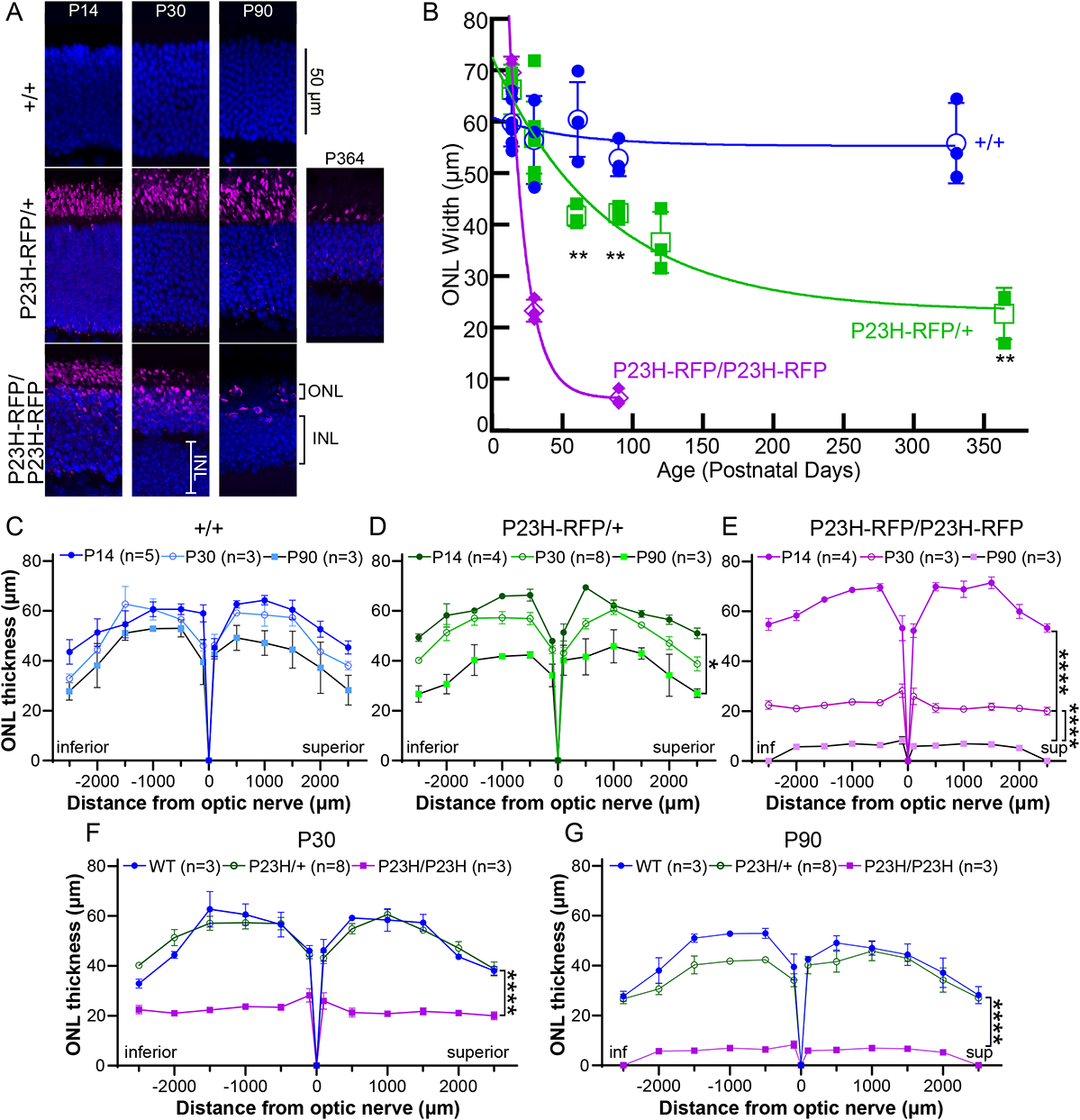
Time course of retinal degeneration in P23H-RFP/+ heterozygous and P23H-RFP/+ homozygous mice. (A) Confocal z-projection images of retinal cryosections from wild-type (+/+), and P23H-RFP heterozygote (het) and P23H-RFP/P23H-RFP homozygous (homo) mice at various time points. DAPI (blue) labels photoreceptor nuclei in the outer nuclear layer (ONL) and the bipolar cell nuclei in the inner nuclear layer (INL). RFP fluorescence (magenta) is primarily located in the inner segment layers of het retinas at all time points, while RFP is also located in the ONL in homo retinas. At P90, the homo ONL is reduced to very few disorganized nuclei surrounded by RFP. (B) Time course plot of ONL thickness between genotypes. Measurements correspond to the ONL thickness of the retina 500 µm inferior to the optic nerve. Unfilled shapes correspond to the mean value, and error bars signify standard error of the mean. Solid lines represent fits to exponential decays to plateau, as described in Methods. Unpaired t-tests were calculated to compare +/+ and het values for significance. Comparisons with significant differences are: P61 +/+ vs P60 het (**p=0.0081), P90 +/+ vs P90 het (**p=0.0072), and P330 +/+ vs P364 het (**p=0.0035). (C) “Spider” plots of ONL thickness for 13 positions in +/+ retinal cryosections from the P14, P30 and P90 time points, spanning positions from the optic nerve to the inferior and superior retina. Positions directly adjacent to the optic nerve position (“0”) correspond to 100 µm superior and inferior to the optic nerve. The most peripheral positions of each plot correspond to 100 µm from the superior and inferior ends of the retina. For each position on the plots, circles correspond to mean values and error bars signify standard error of the mean. (D) Corresponding spider plots for P14, P30, and P90 P23H-RFP/+ het retinal cryosections. Tests for statistical differences between plots from different time points were performed using Two-way ANOVA with Šídák multiple comparisons test. The only test with significance is P14 het vs P90 het (*P=0.0215). (E) Spider plots for P23H-RFP homozygous retinal cryosections at the same time points. Two-way ANOVA with Šídák multiple comparison tests produced significant differences between all time points: P14 vs P30, P14 vs 90, and P30 vs P90 (all ****P<0.0001). (F) Spider plots for all retinal cryosections at timepoint P30 to compare differences between genotypes. Two-way ANOVA with Šídák multiple comparison tests produced significant differences at P30 between: +/+ vs homo and het vs homo. (both ****P<0.0001). (G) Spider plots for timepoint P90 to compare genotypes. Two-way ANOVA with Šídák multiple comparison tests produced significant differences at P90 between: +/+ vs homo. and het. vs homo. (both ****P<0.0001).

At postnatal day 14 (P14) there is no difference in the ONL thickness in the retinas among any of the phenotypes, indicating that the retinas develop normally in the P23H-RFP heterozygous and homozygous mutants (Fig 5A-E). Thereafter, the width of the ONL in the heterozygous retinas declined slowly, approaching a final value of 23 μm with a time constant of 56 days, while the ONL in the homozygous mutants declined much more rapidly, with a time constant of 12 days. By age P90 the ONL in P23H-RFP/P23H-RFP homozygous retinas was reduced to a single, disorganized layer of nuclei (See Fig 5A, 5E, & Fig 5G).

Although the ONL thickness in P23H-RFP/+ heterozygous retinas were significantly reduced in the inferior retina at P60 and P90 (Fig 5B & 5G), the thickness across all regions of the retina in P23H-RFP/+ mutants was not significantly different from that of +/+ retinas at age P90 (Fig 5G). Notably, compared to P14 P23H-RFP/+ measurements, the P90 P23H-RFP/+ ONL thickness was significantly reduced (Fig 5D). By age P364 it was evident that the ONL had been severely reduced in P23H-RFP heterozygous retinas due to nuclei loss (Fig 5B, see example in Fig 5A); the ONL width at 364 days in the heterozygotes was 40% of that in WT at 360 days.

### ERG rod photoreceptor function was moderately diminished in P23H-RFP/+ mice

We next used electroretinogram (ERG) recordings to test and correlate retina visual function to the mild and severe retinal degeneration phenotypes in P23H-RFP/+ mice and P23H-RFP/P23H-RFP mice, respectively. We found generally that ERG waveforms at age P30 correlated to our retinal degeneration phenotypes between the mutant P23H-RFP genotypes compared to +/+ control mice (Fig 6A). In dark-adapted conditions, the scotopic a-wave amplitudes in P23H-RFP/+ mice were significantly reduced compared to +/+ mice at P30 (Fig 6B); however, the a-waves stabilized over time and were not further diminished in P23H-RFP/+ mutants at age P90 compared to P30 (Fig 6C). Scotopic b-wave values were not significantly reduced in P30 P23H-RFP/+ mice compared to +/+ (Fig 6D), and like the a-wave, the b-wave values were not diminished in age P90 P23H/+ mice compared to P30 (Fig 6E). We also measured the implicit times from scotopic ERG recordings, which are times from the a-wave deflection to peak b-wave. In P30 P23H-RFP/+ mice compared to +/+ mice, implicit times were higher at moderate flash intensities, but there was not a significant difference over the entire flash range (Fig 6F).

**Figure 6.**
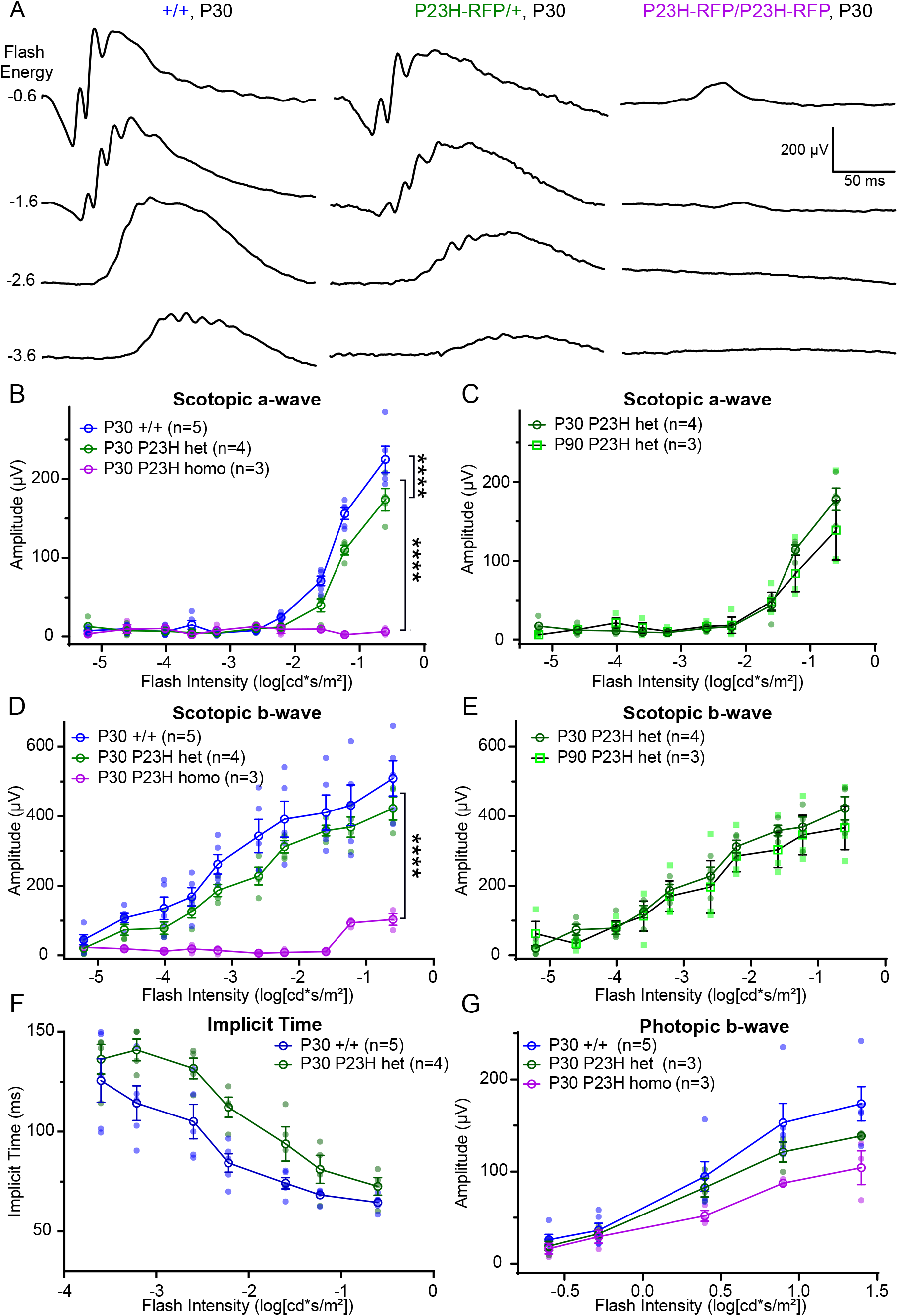
P23H-RFP/+ heterozygous mice have slightly reduced rod photoreceptor electroretinogram (ERG) responses. (A) Example ERG recordings from +/+ wild-type, P23H-RFP/+ heterozygous (het) and P23H-RFP/P23H-RFP homozygous (homo) mice at age P30. (B) Aggregate of P30 scotopic a-wave amplitudes. In all plots, solid shapes are data points, empty shapes signify mean values, and error bars signify standard error of the mean. Statistical comparison tests of all the ERG data were performed using Two-way ANOVA with Šídák multiple comparisons tests. All statistical comparisons of P30 a-wave amplitudes were significant: +/+ vs het (****P<0.0001), +/+ vs homo (****P<0.0001), and het vs homo (****P<0.0001). (C) Scotopic a-wave amplitudes between P30 and P90 P23H/+ het mice were not statistically different (P=0.27). (D) Aggregate of P30 scotopic b-wave amplitudes. Statistically significant differences were calculated for +/+ vs homo (****P<0.0001) and het vs homo (****P<0.0001). WT vs het was not statistically different (P=0.9744) (E) Scotopic b-wave amplitudes between P30 and P90 P23H/+ het mice were not statistically different (P=0.9375). (F) Aggregate implicit times, the time to peak scotopic b-wave post a-wave, in P30 +/+ and het mice. P30 het mice had higher implicit times than +/+ at intermediate flash stages, but the difference over the entire flash range was not statistically significant (P=0.5651). (G) Aggregate of P30 photopic b-wave amplitudes. There were no statistically significant differences among the genotypes: +/+ vs het (P=0.749), +/+ vs homo (P=0.1298), het vs homo (P=0.1553).

P23H-RFP/P23H-RFP homozygous mice have essentially no scotopic ERG response (Fig 6A-B and 6D). The minor b-wave response at high flash intensities in homozygotes could be attributed to cones. We also measured the cone ERG response in light-adapted mice from all genotypes at P30. Photopic b-wave amplitudes were recorded in both P23H-RFP/+ and P23H-RFP/P23H-RFP mice, and although they appeared slightly lower in amplitude, they were not significantly reduced compared to +/+ mice (Fig 6G).

To determine whether there were any visible alterations of cone morphology in the mutants, we used cone arrestin immunofluorescence labeling in our P23H-RFP mice at P30. Cones populated the retinas of P23H-RFP/+ heterozygotes as well as in the retinas of P23H-RFP/P23H-RFP homozygotes despite the massive loss of photoreceptor nuclei in the homozygotes (Fig 7A). At P90 we still observed cones in P23H-RFP/+ heterozygotes, but in P90 homozygotes there were no visible cones remaining, presumably as a result of the nearly complete loss of rods (Fig 7B).

**Figure 7.**
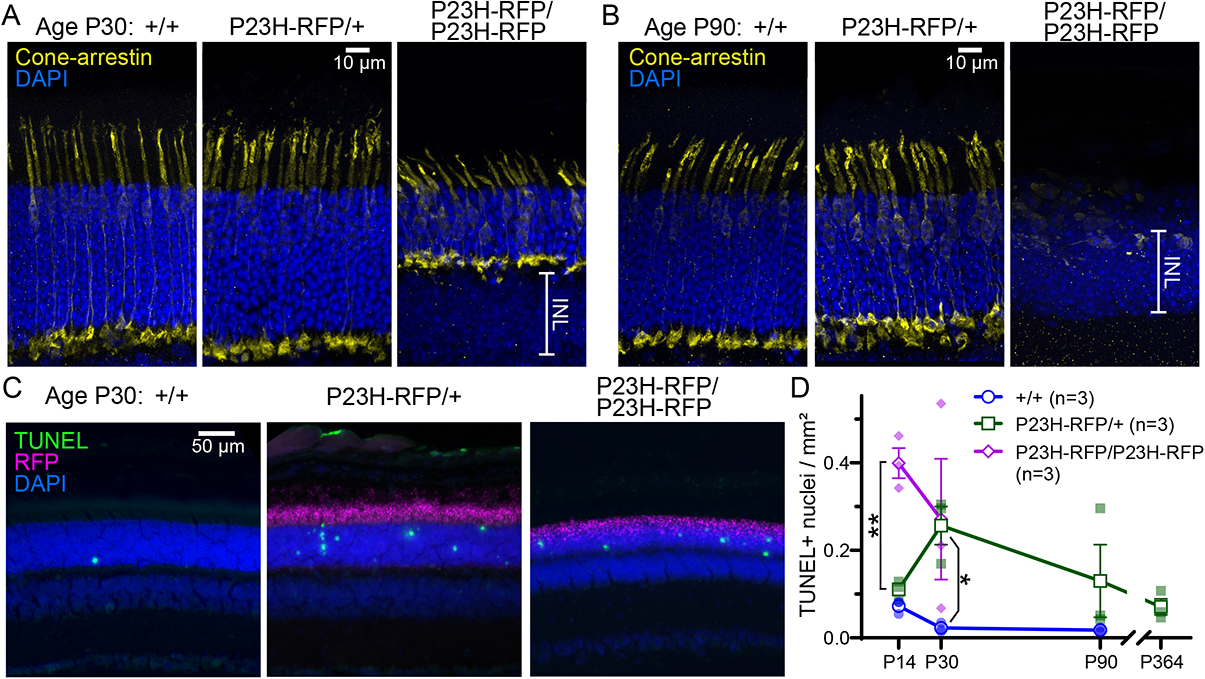
Cone immunolocalization and TUNEL analysis of photoreceptor cell death in P23H-hRho-RFP retinas. (A, B) Examples of cone arrestin immunofluorescence staining (yellow) in retinal sections from ages P30 (A) and P90 (B), among +/+, P23H-RFP/+, and P23H-RFP/P23H-RFP mice. DAPI staining (blue) labels both photoreceptor nuclei in the outer nuclear layer (ONL) and in the inner nuclear layer (INL). (C) TUNEL fluorescence analysis of photoreceptor cell in retinal cryosections. Shown are examples of age P30 retinal sections from +/+, P23H-RFP/+ heterozygous (het), and P23H-RFP/P23H-RFP homozygous (homo) mice with TUNEL+ nuclei (green) within the DAPI stained nuclei of the ONL (blue). RFP fluorescence is magenta. (D) Time course plot of aggregate TUNEL+ nuclei/mm^2^ measurements among all genotypes at multiple time points. Statistical comparisons among groups were performed using unpaired t-tests. At P14, homo retinas have statistically more TUNEL+ nuclei compared to both +/+ (*P=0.0167) and het (**P=0.0098) retinas. At P30, het retinas have statistically more TUNEL+ nuclear compared to +/+ retinas (*P=0.042), but the rate of TUNEL+ nuclei between het and +/+ retinas is not statistically different at age P90 (P=0.312).

To determine the rate of cell death in our P23H-RFP mice, we used terminal deoxynucleotidyl transferase dUTP nick end labeling (TUNEL) fluorescence. We observed TUNEL+ photoreceptor nuclei in the ONL of retinal sections from both P23H-RFP/+ and P23H-RFP/P23H-RFP mice (Fig 7C). We measured the number of TUNEL+ nuclei per mm^2^ area of the ONL at multiple timepoints. At P14, the number of TUNEL+ nuclei in P23H-RFP/+ heterozygous retinal sections were comparable to +/+ sections, while the rate was significantly greater in homozygous retinal sections at P14 (Fig 7D). By age P30, the number of TUNEL+ nuclei was significantly greater in P23H-RFP/+ retina compared to +/+ (Fig 7D); however, at age P90 the TUNEL+ density in heterozygotes was no longer significantly different from that in +/+ mice (Fig 7D).

Overall, we observed that P23H-RFP/+ heterozygous mice have a slow and partial retinal degeneration and mild loss of rod ERG function. At age P90 the heterozygous retinas were still comparable in overall health to control +/+ retinas despite the gross mis-accumulation of P23HhRhoRFP protein in the rod inner segments. The rate of photoreceptor cell death in P23H-RFP/+ retinas indicates that a moderate burst of degeneration begins after P14, which has slowed by age P90. By comparison, the retinal degeneration and ERG phenotypes in homozygous P23H-RFP/P23H-RFP mice were much more severe, such that at P90, nearly all photoreceptor neurons were lost in the retinas of homozygotes.

### Quantification of select mRNA levels from P23H-RFP/+ mouse retinas by Q-RTPCR

As ER stress and the Unfolded Protein Response (UPR) have been proposed to be important in neurodegeneration due to misfolded proteins in photoreceptors (47–51), we examined mRNA levels for several markers of these pathways, using two different “housekeeping” genes for normalization, *RPL19* and *HPRT*. Genes whose messages we quantified included those encoding BiP, CHOP, ATF6, Eif2α, PERK, DRL1 and XBP1. None showed a statistically significant increase relative to both “housekeeping” genes (Fig. 8). We also quantified levels of mRNA transcribed from the *Rhodopsin* locus, using primer pairs that either amplified both human and mouse alleles (mhRho), or the mouse allele only (mRho). Both showed a decrease in message levels in the retinas of heterozygotes relative to those in the WT. The mouse-specific message would be expected to be reduced by ∼50%, based strictly on copy number; however, the reduction was about 70%, whereas the total message derived from both alleles was down about 35% suggesting a down-regulation of both *Rho* mRNAs by 35% relative to WT through either reduced transcription or increased degradation, possibly in response to the presence of aggregated protein.

**Figure 8.**
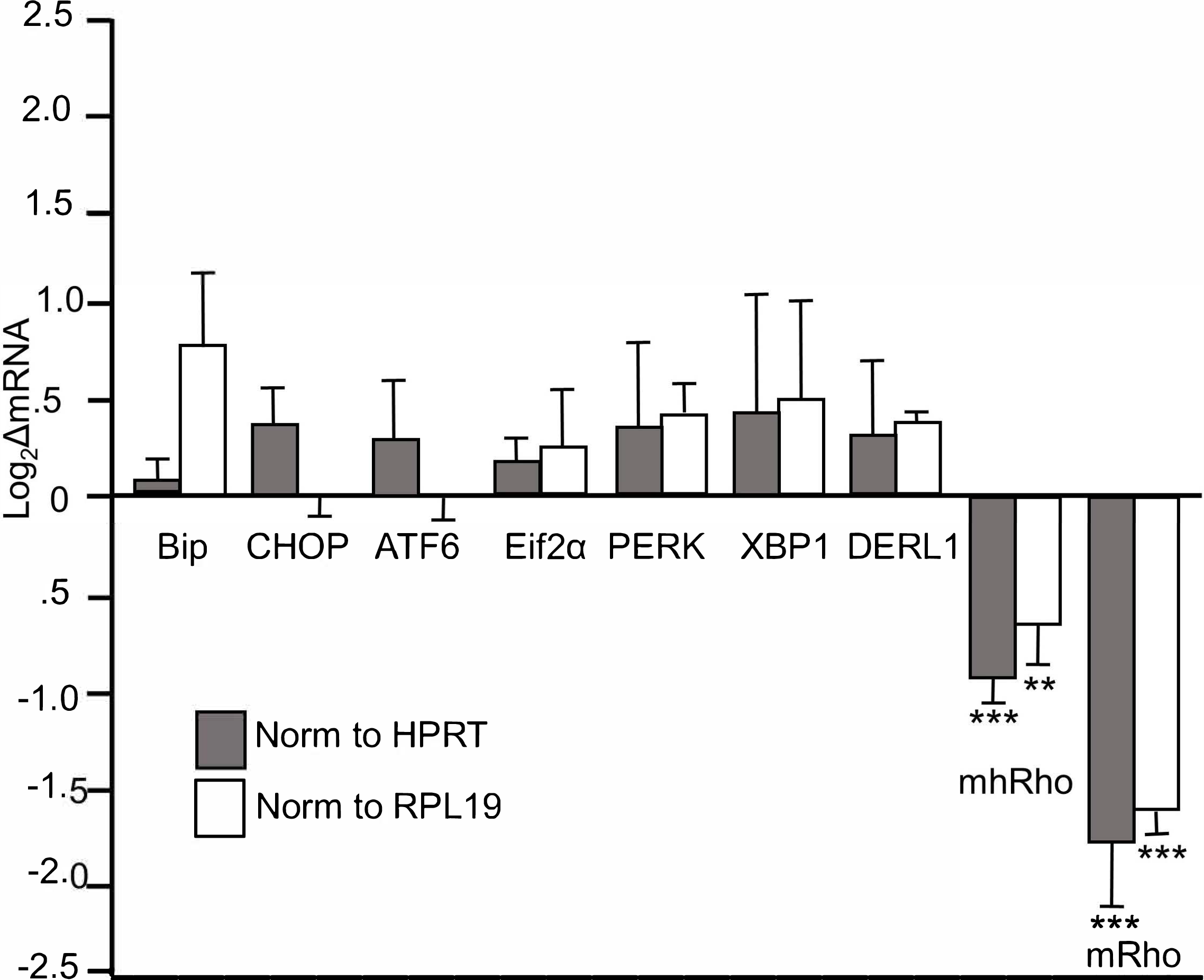
mRNA levels of ER stress and Unfolded Protein Response markers are near normal in P23H-RFP/+ heterozygous retinas at postnatal day 30. Results of Q-RTPCR measurements of the indicated messages in RNA extracted from retinas of heterozygotes as compared to wild-type (+/+) (n=3 for both), normalized to HPRT (dark grey bars) or RPL19 (open bars). Statistical comparisons were made with two-tailed t-tests.: mhRho vs. RPL19 **P = 0.0099; mhRho vs. HPRT ***P=0.000741.; mRho vs. RPL19 ***P=0.000585. mRho vs. HPRT ***P=0.000542.

### P23HhRhoRFP protein was mis-accumulated in the inner segment endoplasmic reticulum

To understand how the mutant P23HhRhoRFP protein is handled by the P23H-RFP/+ rods, we examined the subcellular structures by transmission electron microscopy (TEM) of ultra-thin retina sections following a tannic acid based staining procedure that densely stains internal membranes (8). At age P14, in P23H-RFP/+ heterozygous rods, we observed distinct membranous accumulations in the IS that matched the shape and morphology of the fluorescent RFP+ aggregates (Fig. 9A-B). Compared to +/+ rods, where cytoplasmic membranes are mostly observed in the proximal IS, the membranous accumulations in heterozygous rods were primarily in the form of semi-organized stacks of folded membranes in the distal IS. These membranes often filled the mutant IS cytoplasm, apparently distending the width of the IS itself. Indeed, we measured the average maximum IS width of P14 P23H-RFP/+ rods to be significantly greater compared to the WT average maximum width (Fig 9, see legend). Despite these large aberrant IS membranes in P23H-RFP/+ rods, the morphologies of the BB, CC and OS discs were relatively normal and indistinguishable between P23H-RFP/+ and WT rods at P14, except for the length of the CC, which was significantly longer in mutant rods (Fig 9A-B, see legend).

**Figure 9.**
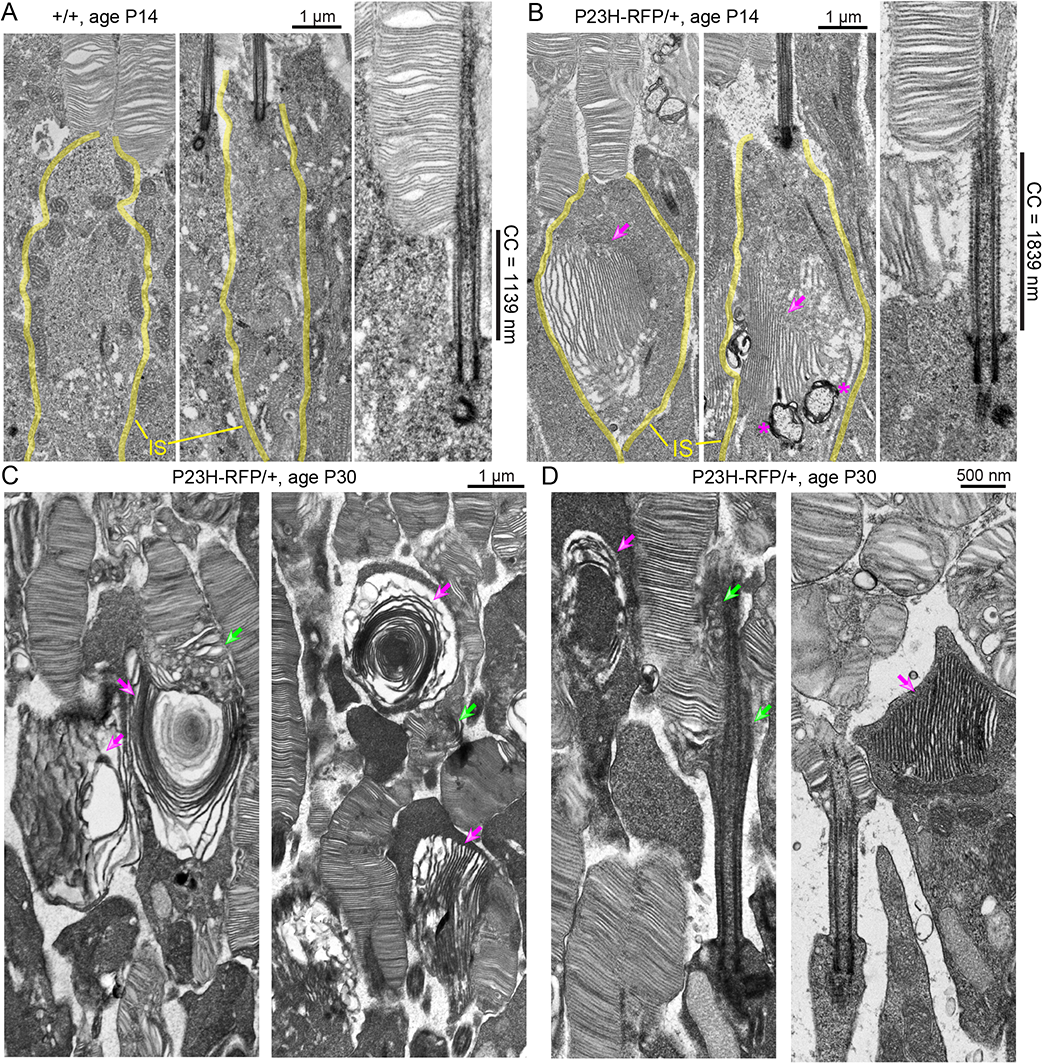
P23H-Rho-RFP/+ mutant rod photoreceptor neurons have distended inner segments filled with ectopic membranes. (A-B) Conventional transmission electron microscopy (TEM) images of rod photoreceptor neurons at age P14 from either (A) +/+ or (B) P23H-RFP/+ heterozygous (het) mice. In each example rod, the inner segment (IS) is outlined with yellow lines. The IS max width in P14 P23H-RFP/+ rods is significantly greater than +/+ rods (het: 2.967 µm ± 0.508 µm (standard deviation, sd) (n=14) vs +/+: 1.974 ± 0.481 µm (sd) (n=19), P < 0.0001, unpaired t-test). Ectopic stacks of IS membranes in the swollen P23H-RFP/+ IS are marked with magenta arrows. Double-membraned autophagy compartments are marked with magenta asterisks in the mutant P23H-RFP/+ IS. In magnified views of example connecting cilia (CC) from each genotype, the length of the CC – measured by the densely stained CC membrane – is indicated. In aggregate, the length of the CC in P23H-RFP/+ rods is significantly greater than +/+ CC (het: 1.548 µm ± 0.206 µm (sd) (n=13) vs +/+: 1.27 µm ± 0.247 µm (sd) (n=11), P = 0.0066, unpaired t-test). (C) At age P30, the ectopic IS membranes in P23H/+ rods appear more dysmorphic compared to P14 (magenta arrows). In addition, some outer segments disc membranes within or adjacent to P23H-RFP/+ rods with IS defects are also disrupted and appear dysmorphic (green arrows). (D) In examples of the CC and basal OS regions of P30 P23H/+ rods, the structure of the CC and basal body remain intact despite being adjacent to ectopic IS membranes (magenta arrows); however, there is evidence that basal OS disc morphogenesis is disrupted possibly due to OS axoneme instability (green arrows).

We also used TEM to examine the morphology of the RFP+ IS aggregates in P23H-RFP/+ rods at P30, where the RFP granule fluorescence appeared less organized than in P14 rods (Fig 4A-B). With TEM, in age P30 P23H-RFP/+ rods, we also observed accumulated IS membranes; however, they did indeed appear more dysmorphic than those at P14, with examples of the membranes wrapping around themselves in whorls (Fig 9C, magenta arrows). As in P14 P23H-RFP/+ retinal sections, the aberrant rod IS membranes at P30 were located among normally formed OS disc stacks, and the CC and BB structures were also morphologically normal at P30 (Fig 9D). Unlike P14 rods, however, we observed some defects in OS membrane morphology at the base of the OS in some P30 P23H-RFP/+ rods. These defects included vesicular and misshapen discs, and an unbound, splayed OS axoneme (Fig 9C-D, green arrows). The results indicate that the mutant P23H-Rho-RFP leads to massive alterations of inner segment membrane structure, accompanied by a least partial disruption of outer segment morphology.

To determine the nature of the membranes in which the RFP fusion protein is located, we used immunofluorescence with antibodies for the endoplasmic reticulum (ER) antigens BiP/GRP78 (binding immunoglobulin protein/glucose-regulated protein 78, an ER lumen chaperone protein) and KDEL (an ER-specific tetrapeptide folding tag), and for the Golgi antigen GM130 (a Golgi-specific membrane marker). In confocal z-projections, we found that BiP ER immunolabeling was co-localized with the RFP+ IS aggregates in rod cells from P14 P23H-RFP/+ retinal sections (Fig 10A). Compared to +/+ BiP labeling, which was largely located in the proximal IS, BiP staining in the mutant P23H-RFP/+ retina was extended into the distal IS and colocalized with the RFP+ aggregates. This result demonstrates that the membrane stacks we observed in the IS of mutant rod via TEM are greatly expanded ER membranes. The Golgi network was unaffected in P14 P23H-RFP/+ retinas and was not co-localized with RFP+ aggregates (Fig 10B).

**Figure 10.**
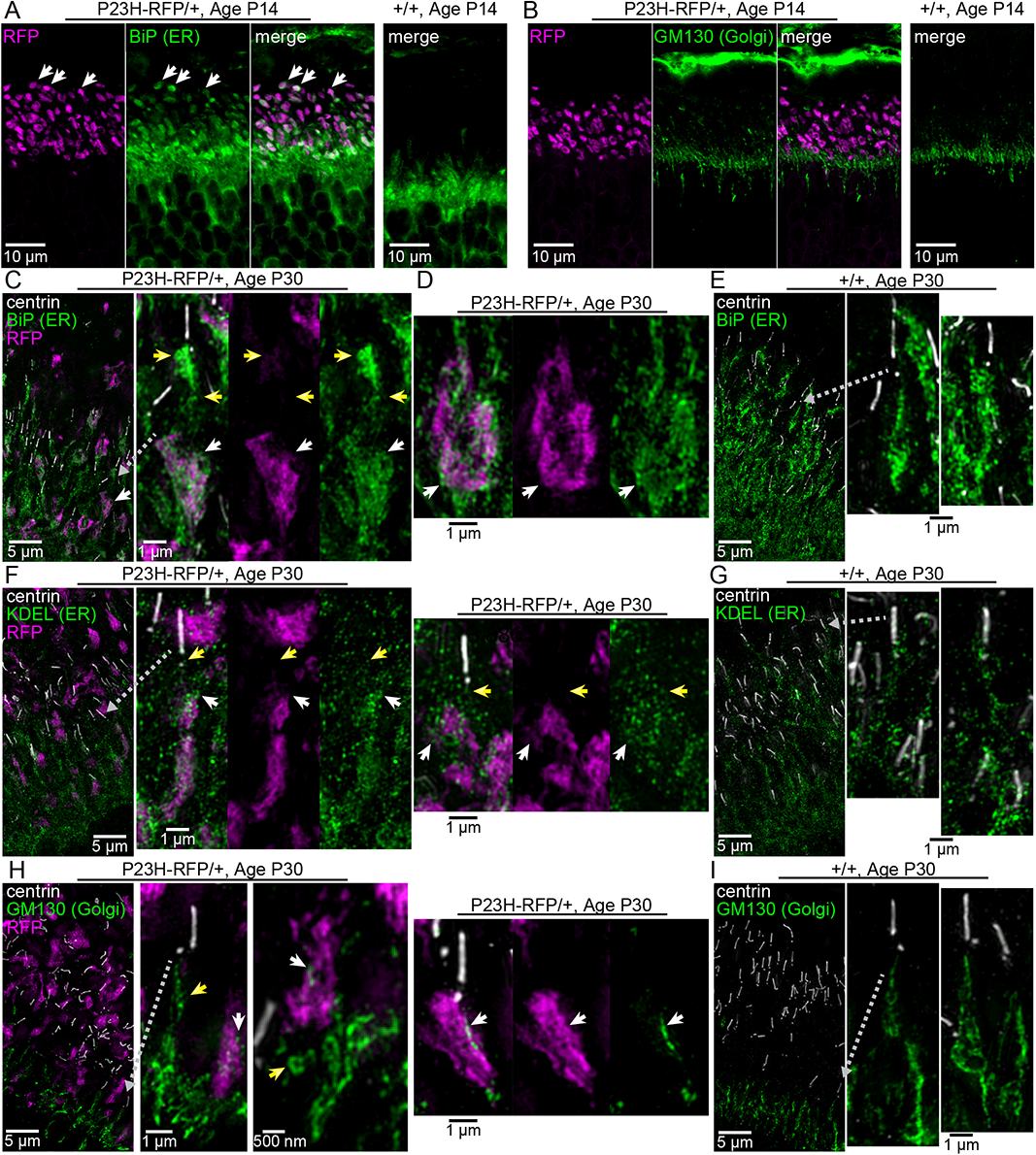
Mutant P23H-hRho-RFP protein is accumulated within ER membranes in P23H-RFP/+ mouse rods. (A-B) Confocal z-projection images of a retinal cryosection from P14 P23H-RFP/+ and +/+ littermate mice, immunolabeled for either the ER lumenal marker BiP/GRP78 (green) (A) or the Golgi marker GM130 (green) (B). P23HhRhoRFP fluorescence is magenta. Co-localized BiP with RFP+ aggregates in the P14 P23H-RFP/+ retina section is marked with white arrows. GM130+ Golgi membranes do not co-localize with RFP+ aggregates and appear unaffected in the P14 P23H-RFP/+ retina compared to +/+ sections. (C) SIM z-projection images of a retina cryosection from an age P30 P23H-RFP/+ mouse that is immunolabeled for BiP (green) and centrin (white), which labels the connecting cilium (CC) and basal body (BB). RFP fluorescence is magenta. In a magnified inner segment (IS), BiP colocalization with a P23HhRhoRFP aggregate is marked with white arrows. BiP+ ER near the cilium that is not colocalized with RFP is marked with yellow arrows. (D) Within the same SIM images, the BiP+ ER tightly surrounds a P23HhRhoRFP aggregate (white arrows). (E) In age P30 +/+ SIM control images, BiP immunolabeling marks the inner segment ER network that leads to the cilium. (F) SIM z-projections of age P30 P23H-RFP/+ retinal sections immunolabeled for KDEL, another ER lumen marker (green), along with centrin (white) and RFP (magenta). KDEL puncta are more diffuse than BiP, but the localization pattern in the IS is similar; KDEL co-localization with the P23H-RFP aggregates (white arrows) and KDEL+ ER near the basal body that is not associated with RFP (yellow arrows) are labeled. (G) In age P30 +/+ SIM control images, KDEL immunolabeling (green) labels puncta throughout the inner segment. (H) SIM images of a P23H-RFP/+ retina cryosection at age P30 immunolabeled for GM130, a Golgi membrane marker (green), along with centrin (white) and RFP (magenta). Overall, The Golgi is proximally localized to the RFP+ aggregates at P30, but in some examples the Golgi network reaches the centrin+ cilium. On a subcellular scale, the Golgi membranes are generally segregated from RFP aggregates (yellow arrows), but small pieces of Golgi membrane are found co-localized with some RFP aggregates (white arrows). (I) In age P30 +/+ SIM control images of retinal cryosections, the Golgi is predominantly dissociated from the cilia; however, examples of +/+ rods with GM130+ Golgi membrane networks that reach the basal body are shown. Gray dotted arrows throughout mark a region that is magnified from the same image.

We used SIM superresolution microscopy to examine the morphology of the ER and Golgi more closely in individual rods in P23H-RFP/+ heterozygous and +/+ retinas at P30. We added centrin immunolabeling to label the CC and BB in these SIM experiments. In P30 P23H-RFP/+ retinas, we again observed BiP co-localization with RFP aggregates in the IS (Fig 10 B-C, white arrows). The BiP+ ER lumen surrounded and was intercalated with the mislocalized P23HhRhoRFP protein that appeared aggregated within the ER membranes. We also observed BiP+ ER in other regions of the IS including at the BB (Fig 10 B-C, yellow arrows). In control wild-type (+/+) P30 retinas, the BiP-positive ER is located throughout the IS in a reticulated morphology that extends to the BB as well (Fig 10E). We observed similar KDEL+ ER localization in P30 P23H/+ rods: both co-localized within RFP+ aggregates (Fig 10F, white arrows) and in the BB region (Fig 10F, yellow arrows); however, KDEL labeling was more punctate than BiP labeling. In P30 +/+ rods, KDEL was also localized throughout the IS and in the BB region (Fig 10G). Although there is much evidence of co-localization of RFP and ER marker signal, the ER markers are not uniformly distributed throughout the clumps of RFP signal, and there are large sections of RFP-positive aggregates without ER marker signal.

We also used SIM to examine the morphology of GM130+ Golgi in P30 P23H-RFP/+ rods. As before we observed the Golgi in the proximal IS of P23H-RFP/+ retinas and segregated from the RFP+ aggregates and the centrin+ CC/BB. In some mutant heterozygous rod IS, however, we observed smaller GM130+ Golgi membranes within the RFP+ aggregates (Fig 10H, white arrows). Also, we found some examples of P30 P23H-RFP/+ rods with more elaborate Golgi that extended into the distal IS and nearby RFP+ aggregates (Fig 10H, yellow arrows). In SIM images of P30+/+ retina immunolabeled for GM130 and centrin, we found most of the Golgi in the proximal IS and dissociated from the centrin+ cilium. Interestingly, however, we did observe some examples of +/+ rods in which the Golgi network was more elaborate and extended up to the BB (Fig 10I).

Finally, in our SIM images we observed both BiP+ and KDEL+ ER localized with the mislocalized P23HhRhoRFP protein in the region of rod synapses in the OPL (outer plexiform layer) of P30 P23H-RFP/+ retinas (Fig 11A-B). GM130-positive Golgi was not localized in the OPL of P30 P23H-RFP/+ retinas (Fig 11C).

**Figure 11.**
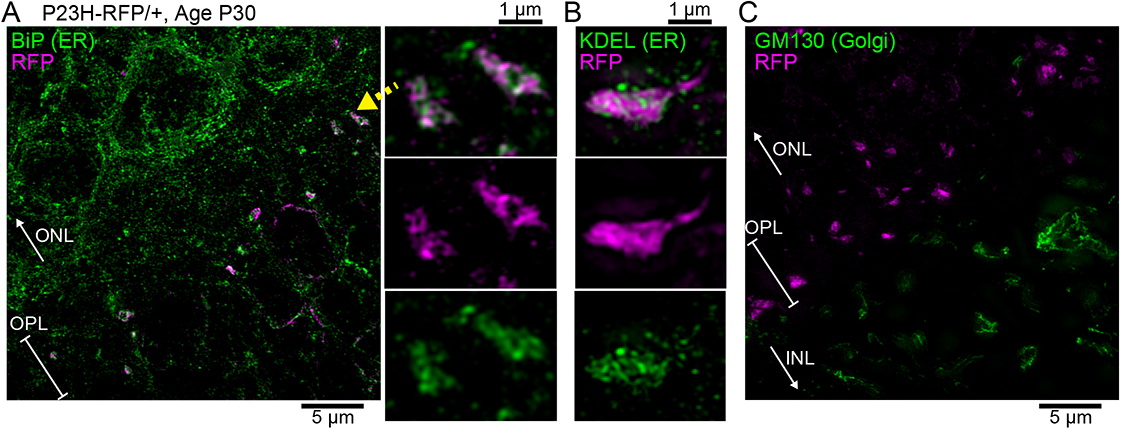
Localization of P23HhRh-RFP protein with the ER in the outer plexiform layer (OPL) of P23H-RFP/+ retinas. (A) A SIM z-projection image of a P23H-RFP/+ retinal cryosection, age P30, immunolabeled for BiP/GRP78 to label the ER (green) in the OPL and surrounding area. P23HRhoRFP protein (magenta) fills the rod photoreceptor synapses in this region. BiP+ ER labeling is present throughout the OPL and the outer nuclear layer (ONL). In a magnified view of a pair of synapses, P23HhRhoRFP protein is co-localized with the BiP+ ER staining. (B) In a similar SIM magnified view of a P30 P23H-RFP/+ retina section immunostained for KDEL (green), P23HRhoRFP protein is also co-localized with KDEL+ ER staining. (C) As a control SIM image, a P30 P23H-RFP/+ retina section is immunostained for GM130 to label Golgi. No Golgi membranes are evident in the OPL; however, they are present in the inner nuclear layer (INL).

## Discussion

The P23HhRhoRFP mouse we introduce here is a potentially useful animal model for adRP that can reveal the subcellular and molecular pathology of the misfolding P23H-Rho mutation on the long-term health of mammalian rod neurons. Tag-RFP-T fluorescence in these mice enables both gross and nanoscopic analysis of P23H-Rho ER accumulation in vivo. The WT-hRho-GFP/P23HhRhoRFP dual color heterozygote demonstrates the different ways the cell processes these two proteins with similar fusions, but with and without the P23H rhodopsin mutation (Fig 3). WT-Rho-GFP fusion protein is restricted almost exclusively to the OS in rods of the Rho-GFP heterozygotes (52), whereas the same fusion construct with a P23H mutation was largely confined to the inner segment and ONL (35), as observed in the new model reported here, consistent with the notion that the P23H mutation is responsible for the disruption in normal trafficking, likely as a result of misfolding (18). Furthermore, with superresolution microscopy, we observed P23HhRhoRFP localization proximal to the connecting cilium and basal body in P23H-RFP/+ heterozygous rods and completely excluded from the OS (Figs 2-3). The exclusive IS mislocalization of the P23H-Rho-RFP protein is unlike P23H-Rho localization in other mouse models, in which there is some detectable OS transport despite other mislocalization and degradation phenotypes (25, 27, 31, 34).

While both P23H-RFP/+ heterozygous and homozygous retinas had mislocalization of P23H-Rho to the IS, rod neuron degeneration in the homozygotes was more severe. In the heterozygotes, despite dramatic P23H-Rho mislocalization and a burst of rod cell death around age P30, the progression of degeneration in P23H-RFP/+ retinas was only partial and relatively slow. By comparison, in the “untagged” P23H-Rho knockin mouse line, heterozygotes lose 43% of their rod population relative to WT by P63 (28); we do not observe such a severe reduction until age P120 (Fig 5B).

The stability of the P23H-RFP/+ heterozygous rods over time enables long term studies in these mice and suggests an adaptation in these rods that provides a neuroprotective effect despite the ER accumulation of P23H-Rho protein. This effect may be related to an ER stress adaptation like the unfolded protein response (UPR), which was originally characterized based on an increase in *BiP*/*GrP78* and *C/EBO protein* (*Chop)* mRNA levels - indicators of an activation of the pancreatic endoplasmic reticulum kinase-like endoplasmic reticulum kinase (PERK) UPR pathway - in transgenic P23H-Rho rats (53). Also in P23H-Rho transgenic rats, overexpression of BiP/Grp78 preserved ERG rod function (54), and overexpression of the BiP-binding ER chaperone ERdj5 preserved photoreceptor survival (55). Although we found no evidence for up-regulation of BiP CHOP or PERK at the mRNA level at P30 in P23H-RFP/+ retinas (Fig. 8), we did not measure corresponding protein levels, or mRNA levels at later ages. We did observe widespread BiP/Grp78 protein in the ER throughout the rod cytoplasm, which co-localized with the ER-accumulated P23H-Rho-RFP protein (Fig. 9).

In contrast, in the untagged P23H-Rho knockin mouse, the inositol-requiring enzyme 1 (IRE1) UPR pathway was shown to be activated concurrent with increased ER-associated protein degradation (ERAD) activity, whereas the PERK pathway was not activated in this model (56). Another study supported a key role for protein degradation by demonstrating that P23H-Rho photoreceptors were preserved by genetically over-activating the proteasome by crossing the untagged P23H-Rho knockin mouse with mice constitutively overexpressing either the PA28α or PSMD11 proteasomal cap protein (57). Therefore, the ER membrane expansion we observed in P23H-RFP/+ rods with TEM (Fig 8) could a be compensatory ER stress mechanism in response to an overload of ERAD and proteasome degradation. Such an adaptation warrants further study. In TEM images of P14 P23H-RFP/+ rods, we also observed double membrane autophagosome-like structures adjacent to the ER (Fig 8A), indicating an autophagy component to the P23H-RFP pathology. Similar double membrane vesiculations were observed in transgenic bovine-P23H-Rho *Xenopus* tadpoles that were exposed to light (23), and more recently LC3-positive autophagosomes were localized adjacent to P23H-Rho protein in the inner segments of these same *Xenopus* rods expressing bovine-P23H-Rho (58).

We observed a deterioration in the morphology of the accumulated ER IS membranes in P23H-RFP/+ rods from age P14 to P30 with both RFP fluorescence (Fig 4) and TEM (Fig 8). In addition to large IS membrane whorls in the IS at P30 (Fig 8C), the ER membrane structures were more disorganized than the tight membrane stacks at P14. We also observed ER at the photoreceptor synapse carrying mislocalized P23HhRhoRFP protein in the OPL of P30 P23H-RFP/+ retinas (Fig 10). Rhodopsin mislocalization to the synapse layer and in the ONL cytoplasm was previously observed in P23H-Rho transgenic mice (31). This suggests that the ER expands throughout the entire cytoplasmic space in rods by P30 as a broad response to the accumulation of misfolded P23H-Rho protein in the ER.

Despite the expanded ER membranes filled with P23HhRhoRFP protein, the morphology of the CC and the OS disc structure was normal in P23H-RFP/+ rods at age P14 (Fig 8A-B). At P30, the OS discs is some P23H-RFP/+ rods were slightly dysmorphic, but the CC structure appeared structurally intact, albeit longer than in +/+ rods (Fig 8C-D). The CC elongation P23H-RFP/+ rods is a surprising result. Such an elongation phenotype was also described in knockout mouse models for 2 CC-localized proteins: male germ cell-associated kinase (Mak) and Huntingtin (59, 60). In both knockouts, the elongated CC was accompanied with aberrant OS morphology and Rho mislocalization. One possible cause of the elongated CC in P23H-RFP/+ mutant rods may be an early imbalance of Rho expression during an early ciliogenesis stage due to mutant P23H-Rho-RFP expression.

Our analysis of the rate of rod neuronal death using TUNEL staining of the ONL in P23H-RFP mouse retinas is additional evidence that P23H-RFP/+ mutant rods undergo a long-term neuroprotective response. Unlike in P23H-RFP/P23H-RFP homozygous retinas where the TUNEL+ rate in the ONL is elevated at P14, the rate of TUNEL+ nuclei spikes at P30 in P23H/+ heterozygous retinas and returns to a level not significantly higher than +/+ retinas by P90 (Fig. 7C). Interestingly, a similar spike in TUNEL staining was also observed in both P23H-Rho transgenic rats at age P18 vs age P30 (32) and in the untagged P23H-Rho knockin mouse at age P19 vs P31 (61).

In conclusion, our RFP fusion knockin model of adRP retinal degeneration caused by the P23H rhodopsin mutation is a unique model for the disease with clear phenotypes that are traceable with both fluorescence microscopy and TEM. The mutant rods in our P23H-RFP/+ mice demonstrate an adaptation that promotes rod photoreceptor survival, and thus the heterozygous mutant retinas have a mild rate of retinal degeneration. As such, our model will be useful to test the variety of proof-of-concept adRP therapies that have been developed across all P23H-Rho models in the field. These include: genetic suppression and replacement strategies (62, 63), CRISPR/Cas9 mutant allele deletions (64–66), genetic inhibition of the autophagy-activating *Atg5* (67), caspase pathway inhibitors (61, 68), a small molecule inhibitor of the photoreceptor specific transcriptional modulator Nr2e3 (69), and pharmacological treatments with valproic acid and other histone deacetylase (HDAC) inhibitors (70). Future studies that investigate the mechanisms of action for these therapies can be tested in this P23H-RFP mouse for the development of the most efficient and synergistic treatments of P23H-Rho and adRP.

## Materials and Methods

### Animals

The P23H-hRho-TagRFP knock-in mice were generated the same way we previously generated P23H-hRho-GFP knock-in mice (35), by gene targeting in the HPRT^−^ embryonic stem (ES) cell line AB2.2 123, which was derived from mouse strain 129SvEv, essentially as described previously (41, 42). The targeting plasmid plasmid contained 5’ and 3’ sequences identical to flanking sequences of the mouse *rhodopsin* gene, an intervening sequence corresponding to the human *Rhodopsin* gene encoding the P23H mutation found in patients, a C-terminal fusion with the fluorescent protein Tag RFP-T, a final C-terminal peptide corresponding to the C-terminal 9 residues of human rhodopsin (TETSQVAPA, the epitope for the 1D4 monoclonal antibody and a putative outer segment targeting signal), a STOP codon, and an endogenous polyadenylation signal, followed by an expression cassette (minigene) for human hypoxanthinephosphoribosyltransferase (HPRT). The HPRT minigene was flanked by loxP sites, so it could be looped out *in vivo* by passing through the germline of GDF-9-iCre mice (44) expressing Cre recombinase in oocytes. The plasmid also contained a minigene for Herpes Simplex Virus thymidine kinase (TK) outside the region of homology for negative selection against non-homologous insertion. We introduced the P23H mutation into the targeting vector by site-directed mutagenesis (QuikChange^Ⓡ^, Stratagene). An ISceI recognition site was engineered into the middle of the first intron in the rhodopsin gene at position 1340 from the start of translation, but it was not used in the experiments described here. The targeting vector was constructed in such a way that there is no Lox site between the MOPS promoter and the rhodopsin transcription unit.

The Darwin Transgenic Core Facility, Baylor College of Medicine, electroporated ES cells and injected correctly targeted ES cells (those selected for HPRT^+^TK^-^ genotype) into blastocysts from albino C57BL/6-Tyr^c-Brd^ mice (43). Founder mice carrying the HPRT-P23H-hRho-TagRFP allele were crossed to GDF-9-iCre mice (44) to remove the HPRT minigene and screened to ensure germline transmission of the correct targeted sequence without HPRT. P23H-hRho-TagRFP mice were extensively backcrossed to C57BL/6 mice. We validated that the knockin was successful by sequencing genomic DNA from the knockin mouse. We verified expression of the P23HhRhoRFP fusion by fluorescence microscopy of retinas and by immunoblotting (Fig. 1, Fig. 2).

The P23H-human-rhodopsin-RFP (P23H-RFP) knockin mice were generated by the BCM Genetically Engineered Mouse core, using a strategy similar to the one previously described for our P23H-Rho-GFP knockin (35). Homologous recombination in ES cells under positive HPRT selection (HAT medium) was carried out with a plasmid containing 5’ and 3’ sequences identical to flanking sequences of the mouse *rhodopsin* gene, an intervening sequence corresponding to the human *Rhodopsin* gene encoding the P23H mutation found in patients, a C-terminal fusion with the fluorescent protein Tag RFP-T, a final C-terminal peptide corresponding to the C-terminal 9 residues of human rhodopsin (TETSQVAPA, the epitope for the 1D4 monoclonal antibody and a putative outer segment targeting signal), a STOP codon, an endogenous polyadenylation signal, followed by an expression cassette (minigene) for human hypoxanthinephosphoribosyltransferase (HPRT). The HPRT minigene was flanked by loxP sites, so it could be looped out in vivo by passing through the germline of a *Zp3Cre* mouse expressing Cre recombinase in oocytes (71, 72).

The junction between the mouse and human *RHO* sequences is at the SacI site in the 5’UTR between the transcription start sites and the translation start sites. Unlike our P23H-Rho-GFP knockin described previously (35), there was no lox site added to the 5’UTR, although there is a lone loxP site remaining in the 3’-end following loop-out of the HPRT minigene. The sequence of the P23H-hRho-TagRFPr allele was verified by Sanger sequencing. We generated a new hRho-EGFP knockin mouse line with an additional C-terminal 1D4 epitope sequence tag by a similar approach, using WT human rhodopsin sequence and EGFP coding sequence instead of Tag-RFP-T. Generation of the original hRho-EGFP knockin line was described previously (41)

Both P23H-RFP and hRho-EGFP-1D4 mice were extensively backcrossed to C57BL/6 (>10 generations). Wild-type (+/+) C57BL/6 littermates were used as controls throughout this study. The following genotyping PCR primers were used for the P23HhRhoRFP knockin allele (5’-GTTCCGGAACTGCATGCTCACCAC) and (5’-GGCGCTGCTCCTGGTGGG), which generate a 975 kb knockin band and 194bp WT band.

All animal research in this study was approved by the Institutional Animal Care and Use Committee at Baylor College of Medicine, and was carried out in accordance with the guidelines set forth in the Statement for the Use of Animals in Ophthalmic and Vision Research of the Association for Research in Vision and Ophthalmology (ARVO).

### Western blotting

Retinal lysates were made by needle extruding mouse retinas in ice-cold Cracking buffer: 25 mM Tris (pH 8), 300 mM sucrose, 15 mM EDTA, 2 mM MgCl_2_ + 1x protease inhibitor cocktail (GenDepot), and lysates were cleared with centrifugation. Protein concentration was calculated with the BCA assay (Bio-Rad), and sample application buffer was added to lysates, which were sonicated to reduce sample viscosity. 100 µg of each lysate sample was loaded on 10% acrylamide gels for SDS-PAGE. Gels were transferred onto nitrocellulose in Tris-Glycine-SDS buffer and probed with primary antibodies: anti-1D4 (Rho) at 1 µg/ml, anti-RFP (Kerafast, 6a11f) at 1 µg/ml, anti-beta-actin (Cell Signaling Technology, 8HD10) diluted 1:1000. Mouse monoclonal anti-1D4 (73) was purified in-house from hybridoma culture medium. Membranes were secondary labeled with one of the following secondary antibodies: anti-mouse or anti-rabbit IRDye680 (LI-COR Biosciences), diluted 1:10,000, and were imaged on a LI-COR Odyssey imager. For clarity, the minimum and maximum input values from Western blot scans were adjusted maintaining a linear slope.

### Retinal immunofluorescence

For cyrosectioning, mouse eyes were enucleated and either 1) cornea punctured and immersion fixed in 4% paraformaldehyde (PFA) diluted in 1x PBS for 45 mins at room temperature before the cornea and lens were removed in 1xPBS, or 2) the cornea and lens were removed and 1xPBS and eye cups were directly embedded Optical Cutting Temperature (OCT) media in plastic cryomolds and flash frozen on a floating liquid nitrogen platform. Fixed eye cups were cryoprotected in 30% sucrose before mounting in OCT in plastic molds and flash freezing. 8 µm – 10 µm cryosections were collected on poly-L-lysine coated glass slides (EMS), and unfixed eye cups sections were immediately fixed with 2% PFA for 2 minutes. This light fixation method was used for all immunolabeling experiments that included anti-centrin cilia immunolabeling. Superior-inferior positions were marked in eyes to be used for retina thickness measurements prior to enucleation to maintain proper orientation throughout fixation and sectioning.

For immunohistochemistry, sections were blocked with either 2% normal goat serum (NGS) (Fitzgerald), 2% bovine serum albumin (BSA) (Sigma), 2% fish skin gelatin (FSG) (Sigma), 0.2% saponin diluted in 1x PBS, or with SUPER block: 15% NGS, 5% BSA, 5% BSA-c (Aurion), 5% FSG, 0.2% saponin in 1xPBS (sections from Fig 7C-D, Fig 9, Fig 10). Blocked sections were probed with 0.5 µg – 2 µg of primary antibodies in the same blocking buffer overnight at room temperature, protected from light. The following antibodies were used: mouse anti-centrin (20H5) (EMD Millipore, 04-1624); rabbit anti-centrin 2 (Proteintech, 15877-1-AP); rabbit anti-BiP (Abcam, ab21685); mouse anti-KDEL (10C3) (Sigma-Aldrich, 420400); mouse anti-GM130 (35/GM130) (BD, 610822); rabbit anti-cone arrestin (EMD Millipore, AB15282).

Sections were washed in 1x PBS and probed with following secondary antibodies: F(ab’)2-goat anti-mouse/anti-rabbit IgG Alexa 488 (Thermo Fisher), F(ab’)2-goat anti-mouse/anti-rabbit IgG Alexa 555 (Thermo Fisher), or F(ab’)2-goat anti-mouse/anti-rabbit IgG Alexa 647 (Thermo Fisher), diluted 1:500 in blocking buffer, for 1 – 1.5 hours at room temperature, protected from light. Sections were counterstained with 0.3 µM DAPI for 1 hour protected from light. Widefield and confocal sections were mounted with #1.5 coverslips in VECTASHIELD (Vector Laboratories), and SIM sections were mounted in ProLong Glass (Thermo Fisher).

Widefield imaging was performed on an inverted Nikon Eclipse TE2000U microscope with mercury lamp excitation, a 10x objective (Nikon, Plan Fluor 10x) and imaging via a Photometrics CoolSnap cf Photometrics digital camera (Roper Scientific) and excitation via a mercury lamp using dichroic mirrors and filters for excitation and emission wavelength selection. Full retina sections were generated by merging overlapping captures in Fiji/ImageJ using the “Stitching” plugin (74, 75). ONL thickness was measured from these full retina section files in Fiji/ImageJ. The position of the optic nerve was designated as position “0”; we then traced the DAPI+ ONL along the superior (positive nm positions) and inferior (negative nm positions) retina. At each position to be measured, we drew a rectangular Region of Interest with edges at the top and bottom of the ONL with a constant perpendicular length of 50 µm at each position and collected ONL thickness measurements.

Confocal imaging was performed on either a Leica TCS-SP5 laser scanning confocal microscope with a 63x oil immersion objective (Leica, HC PL APO CS2 63.0x, numerical aperture 1.40) or a Zeiss LSM710 laser scanning microscopy with a 63x oil immersion objective (Zeiss Plan Apo 63.0x, numerical aperture 1.40). On both systems, sequential imaging scans with 405 diode, 488 nm argon, 543 nm HeNe, and 633 nm HeNe lasers were performed with parameters set to capture sub-saturation fluorescence and to avoid cross-talk. Z-stacks of 1 µm optical confocal sections were projected in Fiji/ImageJ based on maximum intensity values.

SIM imaging was performed on a DeltaVision OMX Blaze Imaging System (v4) (GE Healthcare) with a PLANPON6 60x / NA 1.42 (Olympus) oil immersion objective using oil with a refractive index of 1.520. The system features 488 nm, 568 nm and 647 nm laser lines, and a front illuminated Edge sCMOS (PCO) camera. Images were captured in sequential SIM mode with 15 fringe shift images acquired per optical section per channel. SIM reconstruction was subsequently performed in softWoRx 7 software. Z stacks of 125 nm optical SIM sections were projected in Fiji/ImageJ based on max intensity values.

All images were pseudo-colored and processed for clarity in Fiji/ImageJ; minimum and maximum input values were adjusted maintaining a linear slope. Magnified images of rod cilia were digitally straightened with the Straighten tool in Fiji/ImageJ.

### ERG

Mice were dark adapted overnight and anesthetized with 90 mg/kg ketamine + 14 mg/kg xylazine. 0.5% tropicamide was added as a mydriatic to both eyes with 2.5% phenylephrine hydrochloride and 0.5% proparacaine hydrochloride for analgesia/anesthesia. 2.5% methylcellulose was used to maintain conductivity and for corneal hydration. A ground electrode was inserted into the mouse forehead, and wire electrode loops were placed over each eye. We used the UTAS BigShot Visual Electrodiagnostic System (LKC Technologies) for ERG recordings. Mice were placed within a Ganzfeld chamber and responses were recorded per a sampling rate of 2000 Hz using a 60 Hz notch filter. Scotopic recordings were averaged from 30 flashes for the intensity range of −55 dB to −10 dB at 5 dB increments. Photopic recordings were collected after adapting the mouse to a constant background light (30 cd/m2) for 7 minutes. Recordings were averaged from 60 flashes for −10 dB and 0 dB intensities, and 20 flashes for 5 dB and 10 dB intensities. We converted intensity values from dB to log[cd*s/m²] values based on the instrument’s calibration data.

ERG wave data was visualized and analyzed using a custom Mathematica code previously described (76). Scotopic a-wave amplitude was baseline to the minimum value of the first 35 ms post-flash. Scotopic b-waves were determined after a low pass filtering at 55 Hz; amplitudes were the a-wave minimum to the maximum of the filtered data between 30-120 ms post-flash. Implicit time was the time in ms between the a-wave to the b-wave values. Photopic b-wave waves were also filtered and were baseline to the maximum between 30-120 ms post-flash.

### Transmission Electron Microscopy

The following TEM preparation, based on (8), enhances staining and contrast of internal cell membranes in rod photoreceptors. Mouse eyes for TEM were cornea punctured in the following ice-cold TEM fixative: 2% PFA, 2% glutaraldehyde (EMS), 2.2mM CaCl_2_ diluted in 50 mM MOPS buffer (pH 7.4) and were then immersion fixed in the same fixative for either 1) 45 mins at room temperature with gentle agitation followed by cornea and lens removal and another 1.5 hours on ice, or 2) 4°C overnight before cornea and lens removal. Fixed eye cups were embedded in 4% low melt agarose, and 150 µm vibratome sections were cut along the longitudinal plane. Sections were fixed in 1% tannic acid (EMS), 0.5% saponin (Calbiochem) diluted in 0.1 M HEPES (pH 7.3) for 1 hour at room temperature with gentle rocking. After rinsing again, sections were stained with 1% uranyl acetate (EMS) diluted in 0.1 maleate buffer (pH 6) for 1 hour at room temperature with gentle rocking.

Sections were then dehydrated in the following ethanol series: 50%, 70%, 90%, 100%, 100%, in half dram glass vials filled with 1 mL of dehydrant. For Eponate 12 embedding, sections were additionally dehydrated in 100% acetone 2x, for 15 minutes. Using the solutions of the PELCO Eponate 12 kit with DMP-30 (Ted Pella), a medium hardness resin mix (without accelerator) was prepared and mixed. After dehydrating, the sections were embedded in stages of Eponate 12 resin mix to acetone for the following incubation times: 50%:50% (5 hours), 75%:25% (overnight, 16-20 hours), 100% resin mix (8 hours), 100% resin mix (overnight, 16-20 hours). The DMP-30 accelerator was added to the Eponate 12 resin mix just before mixing or incubating. All embedding steps were incubated on a room temperature roller set to a slow speed.

Resin embedded sections were then mounted in full resin either between two sheets of ACLAR Film (EMS) or in an inverted BEEM embedding capsules (EMS). Mounted sections were cured at 65°C for at least 48 hours. 70-100 nm ultrathin resin sections were cut on a Leica UC6 ultramicrotome with a Diatome diamond knife. Ultrathin sections were collected onto cleaned 100 mesh copper grids (EMS). Grids for TEM were post-stained on glue sticks first in 1.2% uranyl acetate (diluted in Milli-Q water) for 4-6 minutes, then in Sato’s Lead for 4-6 minutes after rinsing in boiled Milli-Q water and drying. Grids were rinsed again and dried before storage. Imaging was performed on either a Hitachi H7500 TEM or JEOL JEM-1400 at magnifications up to 25000x. TEM images were processed for clarity in Fiji/ImageJ by adjusting the minimum and maximum input values (maintaining a linear slope) for contrast and with the Straighten tool.

### RNA extraction and relative quantification of mRNA

Six pairs of retinas from three postnatal day 30 (P30) WT C57BL/6 mice and three P30 P23H-RFP/+ heterozygous mice were collected and homogenized for RNA extraction. Direct-zol™ RNA MiniPrep Kit (Genesee Scientific/Zymo Research, Catalog No: R2051) was used for RNA extraction and 25μl of DNase/RNase free was used for elution. RNA extraction product was diluted 20-fold in DNase/RNase free water (from Zymo Research Direct-zol™ RNA MiniPrep Kit) before recording the UV absorbance spectra on a Hewlett Packard 8452 Diode Array Spectrophotometer. Only RNA samples with 1.7 ≤ *A*_260_/A_280_ ratio ≤ 2.0 was used for Reverse Transcription Polymerase Chain Reaction (RT-PCR) experiments. 5 µl of the RNA extraction product was run on a 1% agarose gel to examine purity. RNA samples with 28S/18S ratio ≥ 2 were used for analysis. The total RNA input for each RT-PCR reaction was 500 ng. RT-PCR was performed using LunaScript RT SuperMix Kit (New England Biolabs – Catalog No: E3010L). Each Complementary DNA (cDNA) sample from RT-PCR was diluted 100-fold prior to quantitative Polymerase Chain Reaction (qPCR). In the Amplifyt™ 96-Well PCR Plate (Thomas Scientific, Catalog No: 1148B05), 4 ul of the diluted cDNA was added per qPCR reaction followed by 16ul of master mix with target gene qPCR primers (See primer nucleotide sequences in Table 1), Luna® Universal qPCR Master Mix (New England Biolabs, Catalog No: M3003L) and DEPC treated water. Per qPCR run, three technical replicates were run per biological replicate. Plates were sealed by LightCycler® 480 Seal Foil (Roche Life Science, Catalog No: 04 729 757 001) then spun down in a mini plate spinner. The qPCR reactions were carried out using the C1000 Touch™ Thermal Cycler (BioRad) and the SYBR fluorescent data collected using the CFX96 Optical Reaction Module for Real-Time PCR Systems (BioRad). Reactions with primers specific for the genes encoding HPRT1 and RPL19 were carried out in parallel for normalization. Data were extracted from CFX BioRad Manager 3.1 as 2007 Excel files. Cq values from 3 technical replicates (with standard deviation ≤ 0.25) were averaged then converted to Relative Quantification Value (RQ value) following Livak Method. Graph design and statistical significance analysis of calculated RQ values were carried out using GraphPad Prism 8.4.3. A 95% confidence interval for a two-tailed t-Test was used as the criterion for statistical significance. *P* values for the two-tailed t-tests are reported.

**Table 1:**
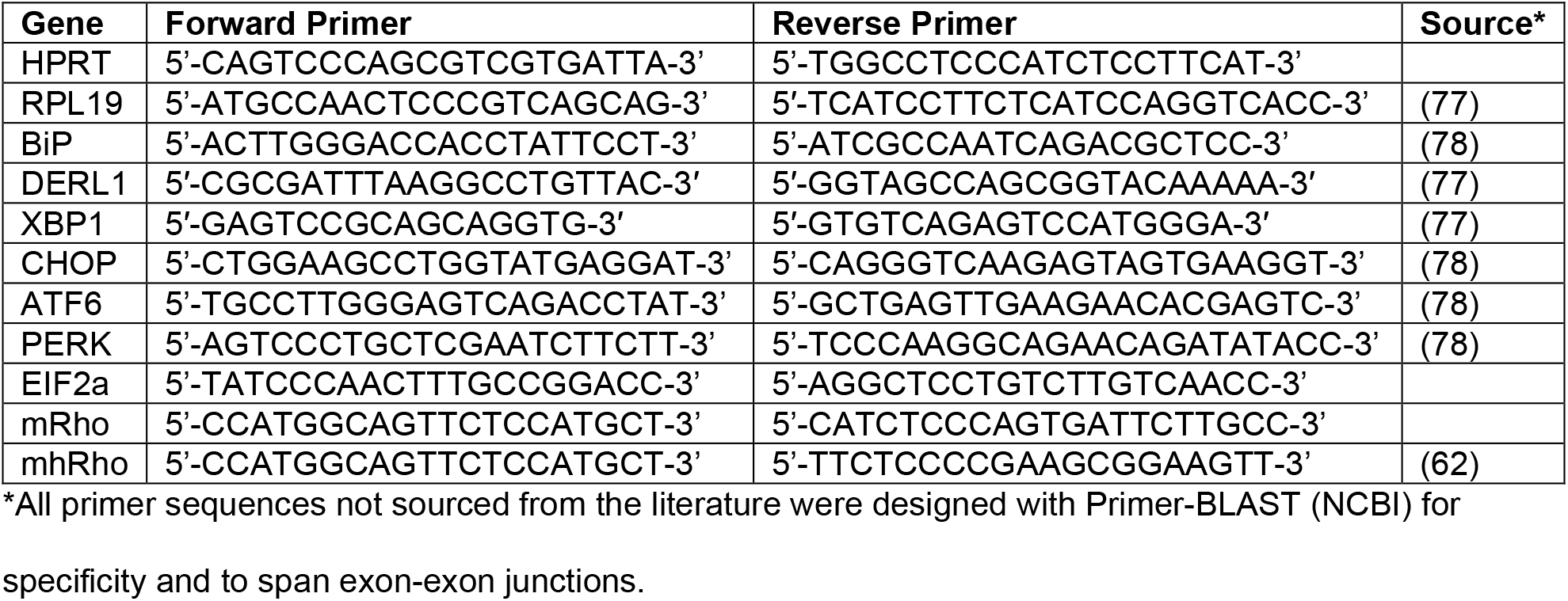
Q-RTPCR Primer Nucleotide Sequences:

### Statistical analysis

Unpaired Student’s t-tests, Two-way ANOVA with Šídák multiple comparison tests, and cubic spline curve fittings were performed in GraphPad Prism® software. Each decline in ONL width with age was fit to a single exponential decay as W(t) = (W(0) – W(∞))exp-(t/τ) + W(∞) Where W(t) is the measured width at age, W(∞) is the plateau value to which it declines, W(0) is an initial value, and τ is the time constant. The data were fit to the equation with W(0), W(∞) and τ as floating fit parameters in a Marquardt-Levenberg least-squares fitting algorithm using Prism®.

## Acknowledgements

The authors would like to thank Dr. Ching-Kang Jason Chen and Dr. Melina Agosto for help with ERG recordings and analysis. This work was supported in part by NIH research grants R01-EY01173, R01EY026545 and R01EY031949, core grants P30EY002520 and P30CA125123, MAR was supported by a grant from the Knights Templar Eye Foundation and NIH grant F32-EY027171.

## Author Contributions

MAR, FC, VN, LK and FH performed experiments, analyzed data and contributed to writing. MAR wrote the original draft. TGW and JHW provided funding, supervised aspects of the projects, analyzed data and edited the manuscript.

